# Transcriptome reallocation during infection reshapes synonymous codon demand in *Botrytis cinerea*

**DOI:** 10.64898/2026.11.04.764316

**Authors:** Jordan A. Dowell, Lori Pradhan, Melanie Madrigal

**Affiliations:** Department of Biological Sciences, Louisiana State University, Baton Rouge, Louisiana 70802, USA; Department of Plant Pathology and Crop Physiology, Louisiana State University AgCenter, Baton Rouge, Louisiana 70802, USA

**Keywords:** *Botrytis cinerea*, codon usage bias, co-transcriptome, translation, transcript allocation, fungal pathogen

## Abstract

While genomic codon usage is fixed within an individual isolate, the synonymous codons represented in its RNA pool can change when transcription shifts among genes. We tested whether this transcript-level phenotype accompanies infection by reanalyzing 39 libraries from three public *Botrytis cinerea* RNA-sequencing (RNA-seq) time series in S. lycopersicum, *Arabidopsis thaliana*, and *Physcomitrium patens*. Gene counts were corrected for coding length, combined with fixed gene-specific codon inventories, and analyzed as centered log ratios within synonymous amino-acid families. Sampling time accounted for 95–98% of fungal codon-demand variation in each series. The association persisted after effective-length correction, unique-only competitive mapping, host-homology exclusion, depth equalization, protein-family weighting, and removal of translation-associated genes. Change vectors over the interpolated, time-aligned 16–24-h interval were positively aligned among studies, whereas later trajectories diverged. Matched reassignments of complete expression trajectories showed more full-series organization than expected in *S. lycopersicum* and *Arabidopsis*, but not in *P. patens*. Codon-preference-weighted abundance and expression breadth also changed through infection, although their directions were not universal. Across systems, large non-translation contributors repeatedly carried ATP-synthase/respiratory and extracellular annotations, although their timing and companion functions differed. Infection-associated transcriptional reallocation therefore repeatedly restructures the synonymous composition of the inferred fungal transcript pool. This establishes a dynamic allocation phenotype and a basis for testing consequences for RNA stability and translation during host–pathogen interactions.

## Introduction

Successful infection requires *Botrytis cinerea* to shift among overlapping biological programs, from germination and penetration to tissue colonization, nutrient acquisition, stress responses and, in some settings, reproduction (Cheung et al. 2020; Wei et al. 2024). Because the genes recruited into these programs differ in both function and synonymous codon composition, transcriptional turnover changes which protein-coding messages are abundant and which codons are represented in the inferred RNA pool. Therefore, while a single isolate’s coding repertoire remains essentially fixed during infection, its RNA allocation among genes is highly plastic. A fixed genome can therefore generate a dynamic transcript-level codon-demand phenotype without changing any individual gene’s coding sequence. Infection-associated differential expression is well characterized, whereas this sequence-compositional consequence of transcriptional turnover has received less direct attention.

Such reorganization is possible because *B. cinerea,* like most organisms, has genes that vary in synonymous codon use. This variation can reflect overlapping effects of mutation and nucleotide composition, genetic drift, gene function, and selection associated with expression or translational decoding (Gupta et al. 2021; Baeza et al. 2024; LaBella 2024; Dahal et al. 2026). For example, codon usage patterns in parasitic fungi can be influenced by directional mutational pressure from the environment and gene expression due to reliance on host metabolic function (Yan et al. 2026). Codon usage can also influence transcript production; in *Neurospora crassa,* synonymous substitutions introduced rare codons that resulted in premature polyadenylation signals, causing early transcription termination and reducing full-length mRNA production (Zhou et al. 2018). While no single statistic captures all these influences, several indices have been developed to describe different patterns of potential codon bias. The effective number of codons (ENC) summarizes how broadly a gene uses synonymous alternatives, with lower values indicating more concentrated usage (Wright 1990). The Codon Adaptation Index (CAI) instead summarizes use of codons enriched in a chosen reference set and is therefore reference-dependent (Sharp and Li 1987). Here, we ask how infection resamples a fixed coding repertoire over time, focusing on within-isolate acclimation of transcript allocation during an infection time course.

Expression weighting turns this idea of potential acclimation of transcript allocation into a dynamic, measurable RNA-level phenotype. Gingold et al. (2012) used expression-weighted codon usage to infer changes in translational demand. Following the same general logic, we define transcriptome-weighted synonymous codon demand as the relative representation of synonymous alternatives implied by RNA abundance. This quantity extends differential-expression analysis by measuring a sequence-compositional consequence of transcriptional reprogramming, but it remains entirely RNA-based. It does not establish ribosome occupancy, translation rate, protein output, or energetic limitation, which require direct assays such as ribosome profiling (Ingolia et al. 2009). However, an RNA-level allocation phenotype may nevertheless be biologically relevant. Cellular decoding capacity can vary with tRNA abundance, charging, wobble pairing, and nucleotide modification (Nedialkova and Leidel 2015; Torrent et al. 2018), while codon identity can influence RNA stability and gene expression in model systems (Presnyak et al. 2015; Zhou et al. 2016). In fungal pathogens, tRNA modification has also been linked to effector production and virulence-related metabolism (Li et al. 2023; He et al. 2024). A transcriptome concentrated in a smaller set of highly expressed genes with high CAI relative to the ribosomal reference set may therefore create a different message environment than one distributed across a broader, more codon-heterogeneous set of metabolic, secretory, stress-response, and host-interaction programs. In addition, nutrient conditions redistribute translational and metabolic capacity in yeast (Metzl-Raz et al. 2017; Yu et al. 2020), and nitrogen availability specifically alters *B. cinerea* infection and virulence-gene expression (Soulié et al. 2020). In *Salmonella sp.,* virulence-associated codon usage enhanced reporter expression relative to ribosomal codon usage under starvation and host-mimicking conditions (Goormaghtigh and Bumann 2026). These observations suggest a resource-allocation hypothesis linking changes in phenotypic functional breadth to changes in transcript-level codon demand.

Evaluating recurrence across infection systems requires comparative and compositional controls. Independent time-series experiments can challenge a pattern observed in one host, but they are not controlled host replicates because host, isolate, tissue, protocol, and developmental state remain partly confounded. We therefore compare direction only across temporally aligned intervals. After normalization within each amino-acid family, synonymous codon shares are compositional by construction: an increase in one is necessarily relative to the others (Aitchison 1986). Within-family log ratios consequently describe relative synonymous representation rather than total amino-acid demand. Finally, matched reassignments preserve complete expression trajectories while breaking their native pairing with coding sequences, allowing us to test whether the observed organization exceeds what transcriptome turnover alone would predict.

We applied this framework to 39 libraries from infection time series in S. lycopersicum, *Arabidopsis thaliana*, and *Physcomitrium patens* (Srivastava et al. 2020; Reboledo et al. 2021; Wei et al. 2024). All fungal reads were represented against the B05.10 coding reference, with RNA-seq sequence concordance used as a sensitivity analysis rather than proof of isolate identity. We predicted recurrent temporal reorganization of synonymous codon demand but not a universal direction, because each infection system redistributes RNA among a different mixture of gene programs. We first asked whether within-family synonymous representation changes with infection time. We then tested which features of that change, including direction and native gene-to-sequence organization, recur across studies. Finally, we examined whether the trajectories coincide with changes in expression breadth, reference-dependent CAI, and source-defined infection programs.

## Materials and methods

### Study design and sequence resources

The primary analysis comprised 39 infection libraries: 12 from *Solanum lycopersicum* (tomato) (PRJNA628162; Srivastava et al. 2020), 18 from *Arabidopsis thaliana* (PRJNA1158549; Wei et al. 2024), and nine from *Physcomitrella patens* (moss) (PRJNA647932; Reboledo et al. 2021; Fig. 1a). The first two studies explicitly used B05.10. The *P. patens* study reported a *B. cinerea* strain isolated from lemon and mapped reads to B05.10 but did not explicitly equate the inoculum with that reference strain (Reboledo et al. 2021). Three deposited 0-HPI libraries associated with PRJNA628162 were synthetic mixtures of mock-inoculated leaf and B05.10 spore RNA and were excluded from infection-time analyses (Srivastava et al. 2020). We analyzed eight libraries from a second *S. lycopersicum* series (PRJNA1056687) separately because only two replicates were available at each time point (Shi et al. 2025). Sample selection, cultivar or accession, tissue, library layout, biological-replicate definitions and exclusions are recorded for all 50 included or explicitly excluded libraries in Online Resource 1, Supplementary Methods SM1 and Online Resource 2: Table S1; study-level statistics are in Online Resource 2: Tables S2–S3.

**Figure 1.**
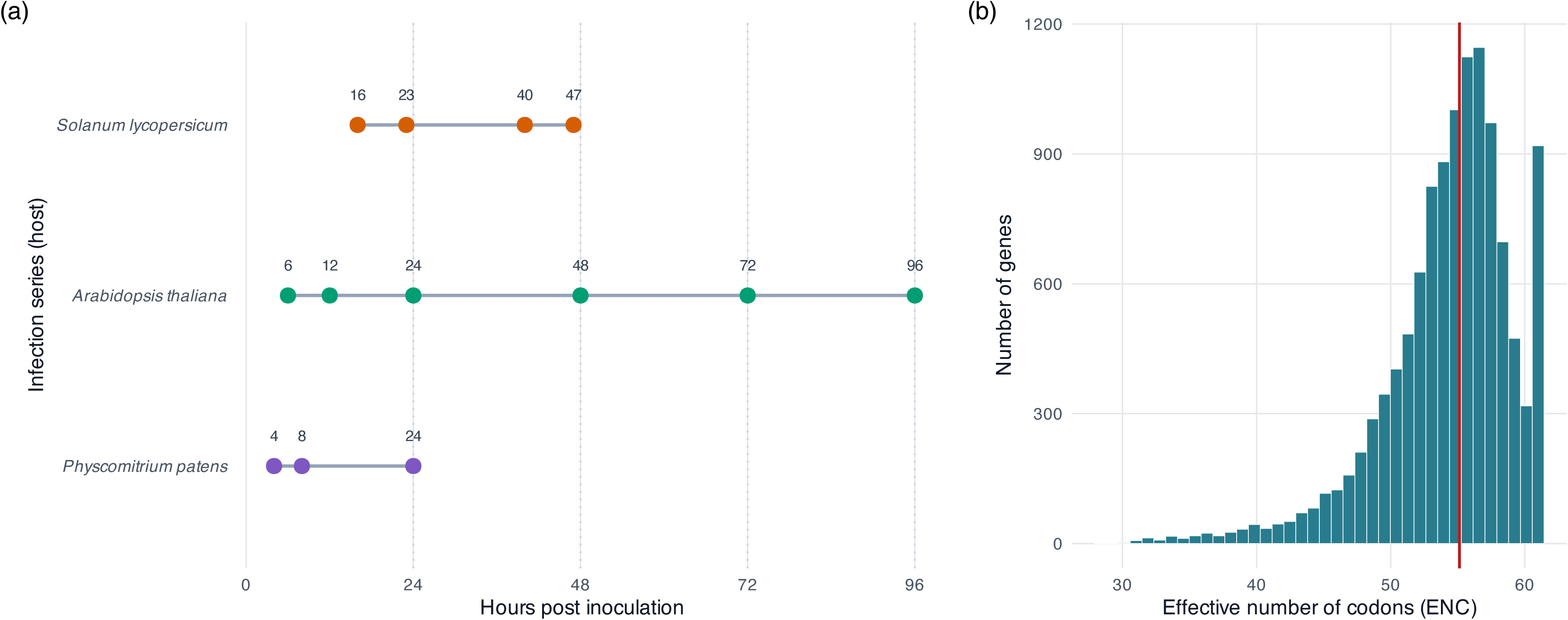
Infection-series sampling and standing B05.10 codon-bias variation. **(a)** Observed infection sampling grids. The primary analysis comprised 39 libraries, with three biological libraries per time point: *Solanum lycopersicum*, PRJNA628162 (16, 23, 40 and 47 HPI; 12 libraries); *Arabidopsis thaliana*, PRJNA1158549 (6, 12, 24, 48, 72 and 96 HPI; 18 libraries); and *Physcomitrium patens*, PRJNA647932 (4, 8 and 24 HPI; nine libraries). Lines connect sampled times only. A second *S. lycopersicum* series, PRJNA1056687 (6, 12, 24 and 48 HPI; two libraries per time point; eight libraries), was used only for supplementary sensitivity analysis and is not plotted. **(b)** Distribution of the effective number of codons (ENC) among 11,698 analyzed B05.10 coding sequences. Lower ENC denotes stronger within-gene synonymous-codon bias. The red line marks the median, 55.13 (interquartile range, 52.15–57.43).

The fungal reference was the gapless B05.10 assembly GCF_000143535.2 (Van Kan et al. 2017). Host coding references were the M82 *S. lycopersicum* gene models (Alonge et al. 2022), TAIR10.1, and *P. patens* V5. Fungal counts for the primary *S. lycopersicum* series used a species-prefixed combined host–fungus genomic reference; host counts for that series and both host and fungal counts for the other three series used competitive host–fungus transcript references. We mapped the supplementary Moneymaker S*. lycopersicum* series against the available M82 transcript reference, and we considered the cultivar/reference mismatch in our interpretation. Gene identifiers were reconciled to 11,698 B05.10 coding sequences (CDSs) passing alignment and coding-quality checks. We used stricter unique-only competitive mappings at mapping-quality (MAPQ) thresholds of 1, 20, and 40; host-only controls where available; whole-gene exclusion of fungal CDSs with host-similar segments; and an expressed-CDS RNA-seq variant panel as assignment, reference-consistency, and isolate-comparability sensitivities. Reference construction and baseline mapping diagnostics are provided in Online Resource 1, Supplementary Methods SM2, and Online Resource 2: Tables S4–S7; length-denominator sensitivity is in Online Resource 2: Tables S8–S11, mapping-environment reconstruction in Online Resource 2: Table S12, strict cross-assignment controls in Online Resource 2: Tables S13–S17 and RNA-seq reference concordance in Online Resource 2: Tables S18–S24.

### Gene-level codon properties and expression-weighted demand

For every fungal coding sequence, we counted sense codons, recorded GC content at synonymous third positions (GC3), and calculated Wright ENC. A ribosomal-reference CAI was calculated from a fixed set of 87 annotated ribosomal genes; the same reference weights were used at every time point and in every fungal study. We therefore interpret temporal CAI results as redistribution of expression among genes with fixed scores, not as changing codon preference. A composition-preserving sequence null randomized synonymous allocation while retaining protein sequence and genome-wide codon totals (Online Resource 1, Supplementary Methods SM3; Online Resource 2: Tables S25–S26).

For each library, we divided raw gene counts by the analyzed coding-sequence length in codons and then normalized to a constant library sum, providing a transcripts-per-million (TPM)-like proxy for relative transcript-molecule abundance. These weights were multiplied by gene-specific codon inventories and summed. This corrects the otherwise duplicated influence of gene length: fragment yield already increases with feature length, while the codon inventory also contains more entries for longer coding sequences. A sensitivity analysis instead used the exact annotation feature length supplied to counting and an effective feature length equal to feature length minus the library-specific mean fragment or read length plus one. Within each amino-acid family, weighted codon totals were converted to proportions and then centered log ratios (CLR) after adding a fixed pseudocount (Aitchison 1986). Only the 59 codons belonging to synonymous families were retained. The response therefore describes relative representation of synonymous alternatives within the same encoded amino acid; it does not report amino-acid demand. Complete definitions and parameter values are given in Online Resource 1, Supplementary Methods SM3–SM4. All analyses used library-level profiles as the units of replication. Time-point means were used for visualization and for defining study-level direction vectors, but they did not increase sample size. Because the data came from different experiments, we normalized within each study and did not use absolute expression abundance for between-study comparisons. Cross-study synthesis compared dimensionless effect sizes, adjusted probabilities, or directions in the common codon alphabet rather than pooling counts across platforms.

### Temporal tests and cross-study comparisons

Within each study, sampling-time separation was tested by permutational multivariate analysis of the 59 CLR coordinates (Anderson 2001). Pseudo-*F* measured separation relative to within-time-point variation, and R² measured the fraction of multivariate variation associated with time. Time labels were permuted 9,999 times, with each library as the experimental unit. Dispersion tests evaluated whether apparent separation instead reflected unequal replicate scatter. Sensitivity analyses equalized leverage among amino-acid families, omitted each library, downsampled counts to the lowest fungal depth within the study, filtered abundance and coding length, downweighted homologous protein families, and excluded either the ribosomal reference or all translation-associated genes (Online Resource 1, Supplementary Methods SM4 and Fig. S1; Online Resource 2: Tables S2 and S25–S41).

Study-centered library profiles and time-point centroids were projected in the common 59-codon space. Directional agreement was summarized by cosine similarity. Comparisons among all three studies used a time-aligned 16–24-HPI contrast obtained by linear interpolation only within each observed range; *S. lycopersicum* and *A. thaliana* were additionally compared over 24–47 HPI. No dataset directly observed both 16 and 24 HPI, and interpolated centroids were never treated as biological replicates (Online Resource 1: Fig. S2; Online Resource 2: Tables S42–S44). We analyzed individual codons using ordinary least-squares models with time point as a categorical predictor. Overall temporal effects used the model *F* test; adjacent-time-point comparisons used pooled-residual-variance *t* contrasts. Benjamini–Hochberg adjustment was applied separately to the 59 overall codon tests and to the complete family of adjacent codon-by-transition contrasts within each study. Fig. 3 displays, descriptively, the 20 largest temporal ranges among the 57 codons significant in all three datasets; symbols show adjusted contrasts with the immediately preceding sampled time point.

### Matched nulls, allocation metrics and host comparisons

To test whether temporal structure followed automatically from changing expression, we reassigned complete abundance trajectories among genes within bins matched for mean abundance, coding length, GC3, and the first two amino-acid-composition principal components. The reassignment preserved all observed transcript trajectories and coding sequences while breaking their gene-specific pairing. A stricter version additionally conditioned assignments on five coarse annotation classes. Full-series pseudo-*F*, time-aligned 16–24-HPI Euclidean displacement, and a trace-standardized organization statistic were compared with matched reassignments. The organization statistic Q was defined as squared displacement divided by the sum of coordinate-wise replicate-estimated contrast variances (Online Resource 1, Supplementary Methods SM5 and Figs. S3–S5; Online Resource 2: Tables S37 and S43–S47).

We calculated transcript-molecule-abundance-weighted mean ribosomal-reference CAI and effective expression breadth (the exponential of Shannon entropy) per library. In the allocation analysis, residual CAI was defined as the residual from a linear model of ribosomal-reference CAI on GC3 and log2 protein length. Genes were assigned to residual-CAI quartiles by genome-wide rank, and library summaries weighted those fixed residuals or quartile memberships by relative transcript-molecule abundance. The secreted/extracellular-gene sensitivity removed the annotated set, renormalized abundance among the remaining genes, and compared the 16–23-HPI and 40–47-HPI groups by exact label reassignment with false-discovery correction. We applied the same length correction, CLR workflow, and matched-trajectory null separately to each host using its own coding reference; we never assigned a shared CAI scale or translation system to host and fungus. To identify functions among large codon-demand contributors, we repeated the adjacent-transition decomposition separately in all three primary series after removing the 456 translation/ribosome-associated genes. Gene-level change vectors were projected onto the total synonymous-demand change, and the top 5% by absolute projection were tested for GO enrichment. For each transition, 4,999 random gene sets preserved coarsened joint 5 × 5 × 4 rank strata of study-wide mean transcript-molecule abundance, analyzed CDS length, and absolute transition-specific abundance change. BH adjustment was calculated within transition, within study, and across all 8,890 tests. Because matching was coarsened, we retain continuous-scale balance diagnostics and interpret enrichment as functional annotation of large RNA-level contributors, not a unique mechanism (Online Resource 1: Supplementary Methods SM6; Online Resource 2: Tables S30, S33 and S35).

Source-defined fungal modules and host-associated gene sets published with the source studies were mapped to the current B05.10 identifiers, with reconciliation coverage audited separately (Online Resource 2: Tables S48–S49). We summarized transcript-allocation trajectories and assessed contributions to first-to-last and 16–24-HPI codon-demand displacement using leave-set-out analysis against 999 matched gene sets per case. The screening matcher prioritized the same coarsened mean-abundance, coding-length, and GC3 cell but expanded to nearby cells, and then to the remaining eligible pool, when exact-cell capacity was insufficient. A stricter sensitivity instead preserved every target cell count exactly and additionally balanced interval-specific absolute RNA-allocation change and amino-acid-composition PC1–PC2; change magnitude was balanced, but signed direction and within-set coordination were not (Online Resource 2: Table S50). Because these tests condition on measured summaries rather than every possible sequence or expression feature, they test disproportionate contribution under their stated nulls rather than unique codon-compositional leverage. Because these programs were partly defined from the same RNA-seq series, same-series applications were explanatory decompositions rather than independent validation.

Additional analyses evaluated *BcNPTII* (Tang et al. 2024) as an external sequence-calibration example, alternative genomic tRNA-decoding assumptions, predefined modification-sensitive codon pairs, and local codon placement near coding-sequence starts. Numbered outputs and their generating analyses are cross-referenced in Online Resource 1, Supplementary Methods SM6–SM9.

Analyses were scripted in R 4.6.0 with fixed seeds. Empirical probabilities used the plus-one correction. Benjamini–Hochberg (BH) adjustment was applied within each declared hypothesis family. Datasets were not pooled at the read-count level. Parameters, seeds, recorded software versions, output manifests, and provenance checks are described in Online Resource 1, Supplementary Methods SM2, SM9, and SM10. OpenAI Codex assisted with analysis scripting and figure formatting, with review by the authors.

## Results

### Infection time restructures synonymous codon demand across all three systems

The common B05.10 reference contributed 11,698 analyzed coding sequences (Fig. 1b). Median ENC was 55.13, consistent with modest genome-wide bias but substantial among-gene heterogeneity (Fig. 1b; Online Resource 2: Table S26). Our primary results establish a nonrandom coding landscape on which temporal transcript allocation can act. In the composition-preserving null, observed median ENC was lower than expected (null mean 56.80), and GC3-adjusted ENC residuals were more variable than expected (observed standard deviation 4.48; null mean 2.67; both empirical *P* < 0.01). Mean length-corrected fungal abundance was positively correlated with ribosomal-reference CAI (observed Spearman ρ = 0.35; null mean 0.02; *P* < 0.01).

Within-amino-acid fungal codon demand differed among time points in all three primary datasets (Fig. 2a, b; Online Resource 2: Tables S2 and S51). *S. lycopersicum* had pseudo-*F* = 52.03, R² = 0.95 and permutation *P* < 0.001; *A. thaliana* had pseudo-*F* = 95.80, R² = 0.98 and *P* < 0.001; and *P. patens* had pseudo-*F* = 76.42, R² = 0.96 and *P* < 0.01. The corresponding BH-adjusted probabilities were all less than 0.01. Fisher’s combined *P* across the three study-level tests was *P <* 0.001. Dispersion was not detectably heterogeneous (*P* = 0.59, 0.29, and 0.56), indicating movement of time-point centroids rather than separation attributable only to unequal replicate scatter. The resultant shift was not attributable to one large synonymous family. Equal family leverage retained significance in *S. lycopersicum*, *A. thaliana,* and *P. patens* (all *P* < 0.01). Dataset-relative early–late contrasts were significant in S. lycopersicum (16–23 versus 40– 47 HPI, *P* < 0.01) and *A. thaliana* (6–24 versus 48–96 HPI, *P* < 0.01), but not in *P. patens* (4–8 versus 24 HPI, *P* = 0.23). All leave-one-library-out tests remained significant (*P* < 0.01), as did all 100 within-study depth-equalization iterations (Online Resource 2: Tables S40–S41).

**Figure 2.**
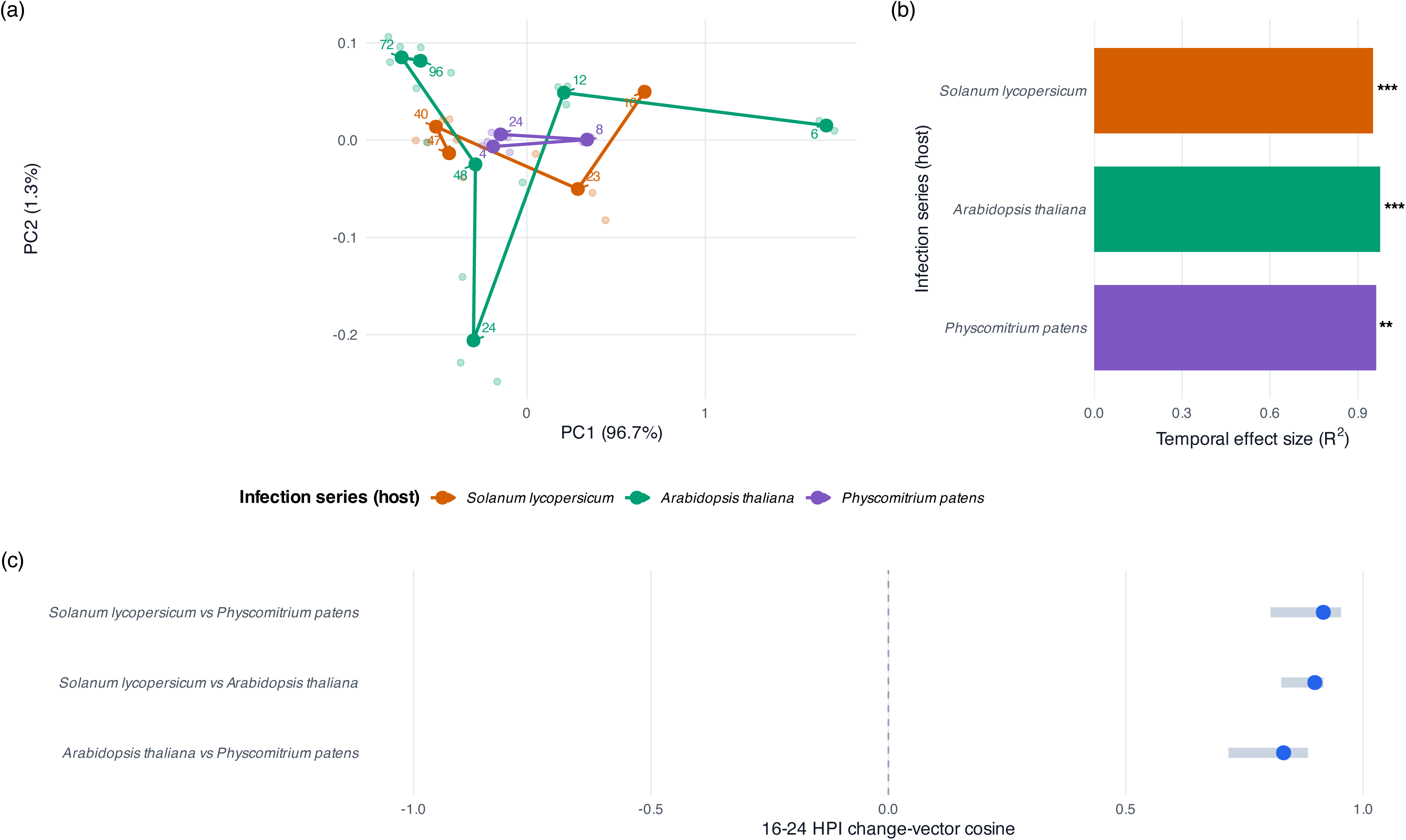
Fungal synonymous-codon demand changes across three host systems. Colors identify the host infection series; all plotted profiles are fungal. **(a)** Principal-component projection of all 39 library profiles after centering each of the 59 within-amino-acid centered-log-ratio (CLR) coordinates within the study. PC1 and PC2 explain 96.7% and 1.3% of the variation, respectively. Faint points are biological libraries; larger points are time-point centroids, arrows show sampling order, and numeric labels give HPI. **(b)** Within-study temporal R² from the canonical 9,999-label-permutation analysis. Stars denote BH-adjusted probabilities across the three study-level tests: ***, *q* < 0.001; **, *q* < 0.01; *, *q* < 0.05. R² values summarize different time grids and are not formal between-study comparisons. **(c)** Pairwise cosine similarity of fungal change vectors over the time-aligned 16–24-HPI interval: *S. lycopersicum–Arabidopsis*, 0.90 (95% bootstrap CI 0.83–0.92); *S. lycopersicum–P. patens*, 0.92 (0.80–0.95); and *Arabidopsis–P. patens*, 0.83 (0.72–0.88). A cosine of 1 indicates parallel change, 0 orthogonal change, and −1 opposite change.

Abundance filters, short-CDS exclusion, and inverse protein-family-size weighting retained every fungal temporal effect (Online Resource 1: Fig. S1; Online Resource 2: Table S27). Excluding 87 ribosomal-reference genes or all 456 ribosome/translation-associated genes also retained the trajectories. Following broad translation-gene exclusion, R² was 0.91, 0.95, and 0.96, and each sensitivity profile correlated at least 0.99 with its corresponding primary profile. Thus, depth equalization and the tested gene-set sensitivities do not explain the result. However, they do not exclude low-level host-to-fungus cross-assignment. Replacing analyzed CDS length with the exact feature length and then with a library-specific effective feature length retained the temporal effect (Online Resource 1: Fig. S6; Online Resource 2: Tables S8–S10). Under the effective-length definition, R² was 0.95, 0.97, and 0.95 for S. lycopersicum, *A. thaliana*, and *P. patens*, with all BH *q* ≤ 0.01. Even there, the effective-length and primary distance matrices correlated at 0.9997, and their first-to-last change vectors had a cosine of 0.998. The corresponding distance correlations were 0.998 and 0.995 in *A. thaliana* and *P. patens*, and direction cosines were 0.998 and 0.97. This sensitivity corrects feature-end opportunity while holding alignments and assignments fixed. Stricter competitive remapping also retained the temporal pattern (Online Resource 1: Fig. S7a; Online Resource 2: Tables S13–S14). With unique alignments at MAPQ ≥20, R² was 0.95, 0.98, and 0.96 for *S. lycopersicum*, *A. thaliana*, and *P. patens*; at MAPQ ≥40, it was 0.95, 0.98, and 0.96. All six tests remained significant (*P* ≤ 0.01), and every strict profile correlated at least 0.99 with its study’s primary CLR profile.

Median fungal-assignment retention at MAPQ ≥20/40 was 95.9%/77.9% in S. lycopersicum, 99.2%/93.0% in *A. thaliana*, and 99.2%/92.7% in *P. patens*. The primary ≥90%-identity/50-nt host-homology screen flagged only 44, 29, and 20 fungal genes, respectively; excluding those genes changed R² by less than 0.0005 under the primary counts.

Host-only controls provided additional context (Online Resource 1: Fig. S7b; Online Resource 2: Tables S15–S16 and S52). The three uninfected *A. thaliana* libraries produced 0.63–10.55 apparent fungal assignments per million host-assigned fragments at MAPQ ≥20. The 4-and 24-HPI P. patens mocks gave 0.45 and 0.53 per million, but the 8-HPI mock was anomalous at 311.50 per million and remained elevated at MAPQ ≥40 (292.42 per million). Only one of its 1,618 MAPQ ≥40 assignments overlapped a fungal gene in the primary host-homology mask.

Removing every fungal gene with any MAPQ ≥20 mock signal (90 genes in *A. thaliana* and 1,045 in *P. patens)* retained temporal R² of 0.98 (*P* < 0.001) and 0.96 (*P* < 0.01); less expansive total-count and recurrent-control blacklists also retained significance. These results argue against generic host-sequence cross-mapping as the source of the temporal effect, but the anomalous *P. patens* control signal remains unexplained. No equivalent pure-host *S. lycopersicum* control was available because the deposited 0-HPI libraries contain B05.10 spore RNA.

The expressed-CDS concordance panel clarified the isolate claim (Online Resource 1: Fig. S8; Online Resource 2: Tables S18–S23). Robust callable sites covered 21.8%, 16.0% and 7.0% of the analyzed B05.10 CDS reference in *S. lycopersicum*, *A. thaliana* and P*. patens*, respectively. Among 3,755,363 *S. lycopersicum* and 2,757,262 *A. thaliana* positions meeting the stringent pooled-depth, allele-fraction, and replicate-support criteria, each study had one nonreference consensus call (0.27 and 0.36 calls per Mb; reference concordance >99.99%). This is consistent with the reported B05.10 identity of those studies at callable expressed-CDS positions. *P. patens* had 1,043 nonreference calls among 1,211,011 callable positions, distributed among 512 genes (861.3 per Mb; 99.9139% reference concordance); 926 calls were synonymous, and 117 were nonsynonymous under the B05.10 coding model. At 829,512 positions callable in all three studies, all consensuses agreed at 99.91%; 707 positions conflicted, and no common site carried the same alternate allele in all three. The *P. patens* reads are therefore highly compatible with the B05.10 coding reference but are not sequence-identical to it, consistent with the source paper’s description of a lemon-derived isolate.

Applying all 1,043 supported *P. patens* substitutions to the affected coding sequences did not materially change the *P. patens* result (Online Resource 2: Table S24). Temporal R² was 0.96 with the B05.10 inventory and 0.96 with the partially corrected inventory; both gave permutation *P* < 0.01 under the same 9,999 label permutations. Reference and corrected CLR profiles correlated at 0.99, their distance matrices at 0.99, and their first-to-last vectors had a cosine of 0.99. This indicates that supported differences at callable expressed-CDS positions do not materially account for the broad *P. patens* temporal trajectory. The panel and correction remain reference-mapped, leave non-callable positions as B05.10, and cannot replace whole-genome isolate authentication.

### A shared early component gives way to study-specific trajectories

The three study-centered centroid paths differed in shape, reflecting their distinct sampling grids (Fig. 2a; Online Resource 2: Table S53). In the time-aligned 16–24-HPI contrast, fungal change vectors were positively aligned: cosine similarity was 0.90 for S. lycopersicum–*A. thaliana* (library-bootstrap 95% confidence interval [CI] 0.83–0.92), 0.92 for *S. lycopersicum–P. patens* (0.81–0.95) and 0.83 for A. thaliana –P. patens (0.72–0.88; Fig. 2c; Online Resource 1: Fig. S9; Online Resource 2: Table S42). *S. lycopersicum* and *A. thaliana* were nearly orthogonal over 24–47 HPI (cosine = 0.05; point estimate only; Online Resource 2: Table S42). Thus, recurrence extended to early direction under this time-aligned sensitivity analysis, but complete or non-time-aligned trajectories were not treated as equivalent.

The second *S. lycopersicum* study further bounded directional replication. It showed a large temporal-separation estimate (R² = 0.87) but did not pass the 9,999-permutation test after length correction (*P* = 0.056). Family-balanced analysis was marginal (*P* < 0.05), the dataset-relative early–late contrast was not significant (*P* = 0.11), one omission test was nonsignificant, and only expression breadth (not weighted CAI) varied detectably across time (*P* < 0.01 and *P* = 0.058, respectively). Its matched 16–47-HPI vector also opposed the primary *S. lycopersicum* trajectory (cosine = −0.964; Online Resource 1: Fig. S10; Online Resource 2: Tables S3 and S54). Two libraries per time point precluded a reliable dispersion test, and we mapped its Moneymaker reads against M82 transcripts.

At the coordinate level, 57, 59 and 59 synonymous codons showed significant overall temporal effects in *S. lycopersicum*, *A. thaliana* and *P. patens*, respectively (Online Resource 2: Table S55). Fig. 3 displays the 20 largest temporal ranges among the 57 codons significant in all three datasets; the selected coordinates changed at different times and were not uniformly monotonic. This combination is consistent with a shared component in the early point estimates superimposed on dataset-specific trajectories. Importantly, CLR coordinates within each synonymous family sum to zero, and transcriptome reallocation affects many genes simultaneously. The codon-level tests localize the multivariate time effect and show that it is distributed broadly rather than confined to one or two unusual codons.

**Figure 3.**
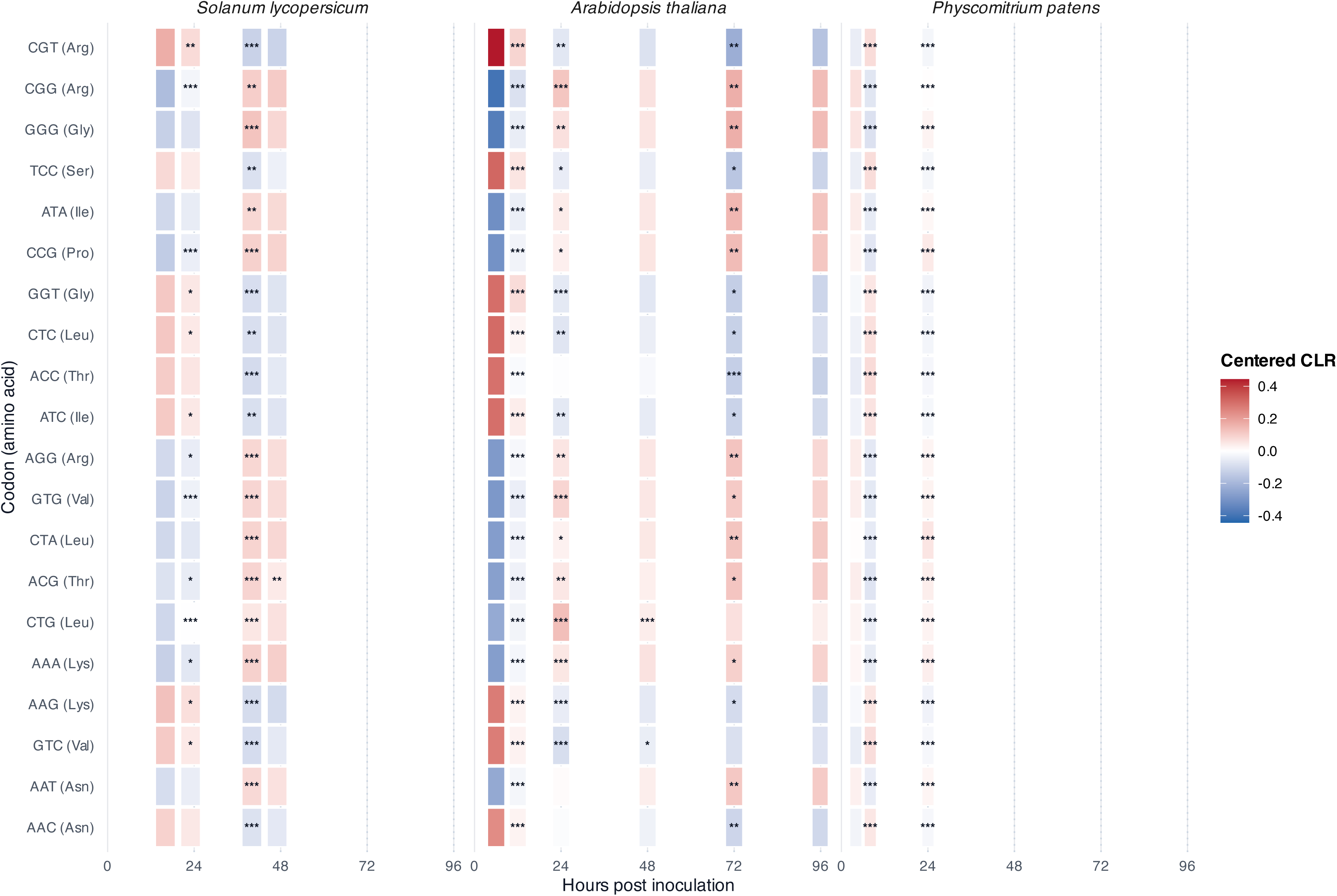
Significant codons follow distinct temporal trajectories across host systems. Cells show each codon’s mean within-amino-acid CLR demand at a sampled time, centered on that codon’s temporal mean within study; blue and red therefore indicate lower and higher representation, respectively, relative to the same codon’s study-specific temporal mean. Each cell summarizes three biological libraries. The figure displays the 20 largest temporal ranges among the 57 codons with significant overall time effects in all three datasets; this ranking is descriptive. Stars compare a time point with the immediately preceding sampled time after BH correction across all codon-by-transition contrasts within that study: ***, *q* < 0.001; **, *q* < 0.01; *, *q* < 0.05; no symbol, *q* ≥ 0.05 or the first sampled time. Panels share a 0–96-HPI axis and dotted guides mark 24-h intervals, but adjacent intervals differ in duration. Host names identify infection series; every plotted quantity is fungal.

### Native gene-to-sequence pairing contributes differently by system and interval

Matched trajectory reassignment distinguished temporal movement from gene-specific codon organization (Fig. 4a; Online Resource 1: Fig. S3; Online Resource 2: Table S37). Full-series fungal pseudo-*F* exceeded the standard abundance/sequence-matched null in *S. lycopersicum* (observed 52.03; null 95% interval 12.76–33.70; empirical *P* < 0.001) and *A. thaliana* (95.80; 19.69–41.68; *P* < 0.001), but not in *P. patens* (76.42; 28.87–93.97; *P* = 0.10).

**Figure 4.**
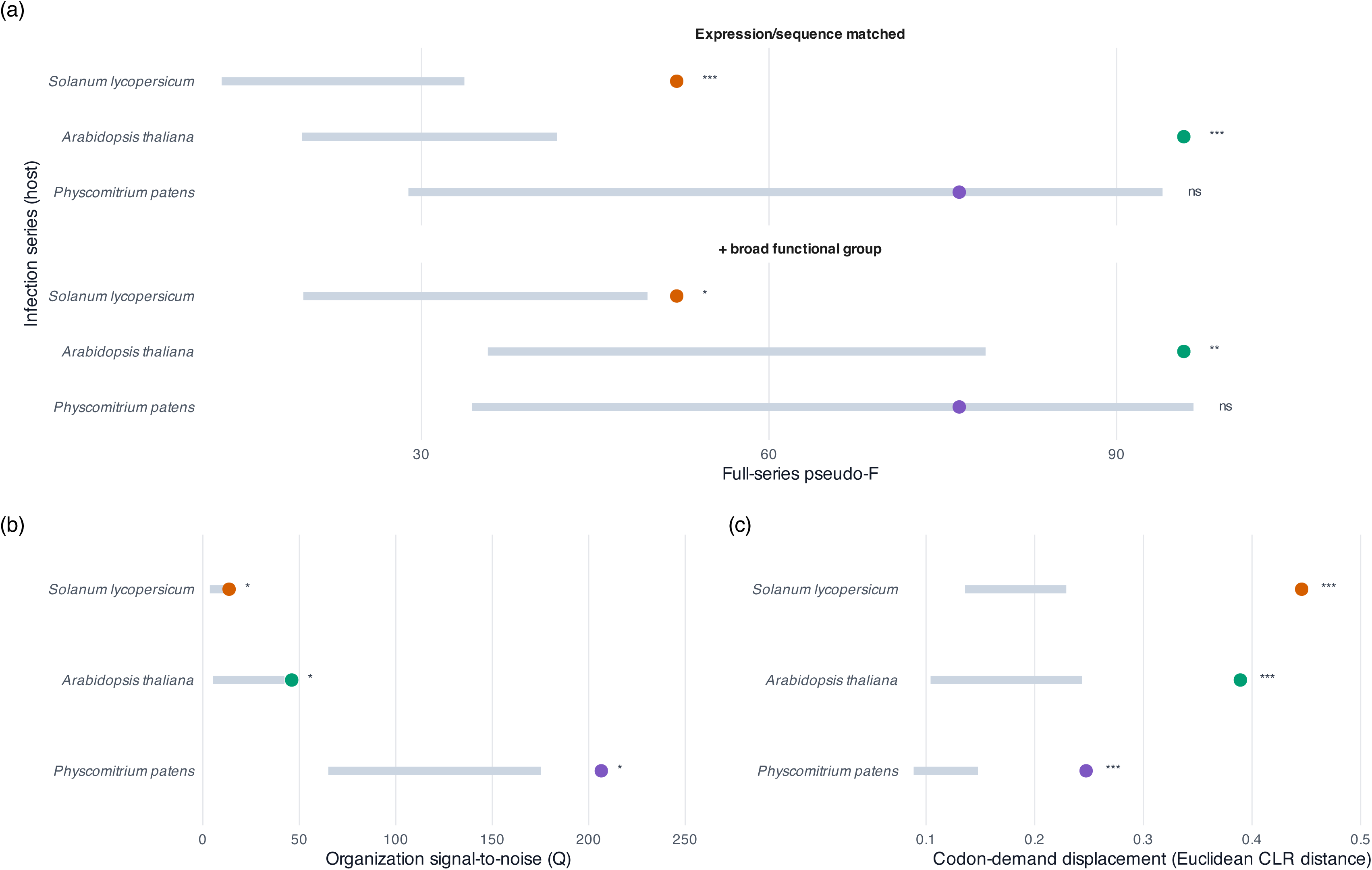
Matched trajectory reassignments distinguish temporal movement from native gene-to-sequence organization. Points are observed fungal statistics; grey segments delimit the 2.5th–97.5th percentiles of the matched-null distribution, and symbols report BH-adjusted probabilities: ***, *q* < 0.001; **, *q* < 0.01; *, *q* < 0.05; ns, not significant. **(a)** Full-series pseudo-*F* compared with an expression/sequence-matched reassignment null (9,999 reassignments) and a second null additionally conditioned on five broad functional classes (1,999 reassignments). Expression/sequence-matched *q* values were 0.0006, 0.0003, and 0.10 for *S. lycopersicum*, *A. thaliana*, and *P. patens*; function-conditioned values were 0.02, 0.003, and 0.15. **(b)** Organization statistic Q for the time-aligned 16–24-HPI contrast, defined as squared CLR displacement divided by the sum of coordinate-wise replicate-estimated contrast variances; *q* = 0.02 in every system. **(c)** Euclidean CLR displacement over the same contrast; *q* = 0.0005 in every system. Panels b and c use the function-conditioned null.

Conditioning reassignments on five coarse functional classes retained full-series excess in S. lycopersicum and *A. thaliana* (all BH *q* < 0.05) but not *P. patens* (*q* = 0.15; Online Resource 1: Fig. S5; Online Resource 2: Tables S46 and S47). Conditioning on these broad classes therefore did not remove the two longer full-series signals and does not rule out finer pathway-level turnover. The time-aligned 16–24-HPI comparison produced a complementary pattern (Fig. 4b, c; Online Resource 1: Figs. S4–S5; Online Resource 2: Tables S43–S46). Euclidean CLR displacement exceeded both matched nulls in all three datasets (function-conditioned BH *q* < 0.001 for each). Trace-standardized organization Q also exceeded the function-conditioned null in all three (*q* < 0.05 for each). Under the standard full-series matching scheme, Q evidence differed among datasets (*q* = 0.09, 0.04 and 0.003 for *S. lycopersicum*, *A. thaliana* and *P. patens*). Thus, the early result is sensitive to the null’s matching definition for *S. lycopersicum,* while raw movement is consistently greater than expected.

### Transcript allocation broadens without a universal codon-preference trajectory

Length-corrected, abundance-weighted ribosomal-reference CAI varied with time in *S. lycopersicum*, *A. thaliana* and *P. patens* (all BH *q* < 0.05; Fig. 5a; Online Resource 1: Fig. S11; Online Resource 2: Tables S56–S57). Effective expression breadth also varied in every dataset (all *q* < 0.05; Fig. 5b). *S. lycopersicum* and *A. thaliana* ended their sampled intervals with lower weighted CAI and broader expression than at their first time points. *P. patens* instead showed a transient CAI maximum at 8 HPI while breadth increased by 24 HPI. The recurrent result was therefore redistribution of preference and breadth, not a universal monotonic trajectory or a tested correlation between the two metrics.

**Figure 5.**
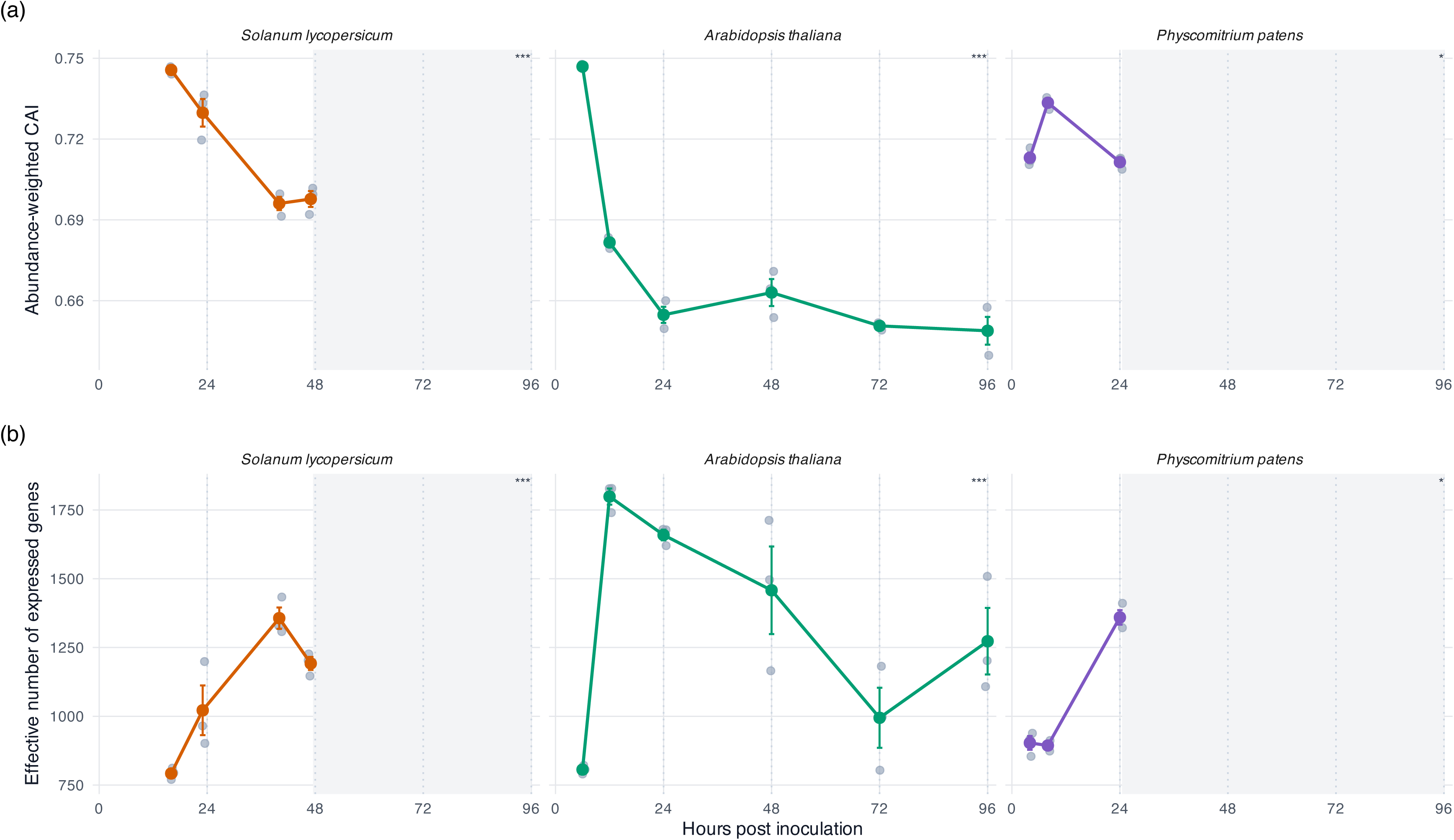
Fungal transcript allocation changes across host systems. Grey points are libraries; colored points and lines are time-point means, bars are ± standard error, and grey shading denotes time beyond a dataset’s final observation. **(a)** Relative-transcript-molecule-abundance-weighted ribosomal-reference codon adaptation index (CAI). BH-adjusted *q* values were < 0.001, < 0.001, and 0.02 for *S. lycopersicum, A. thaliana,* and *P. patens*. The gene-level CAI score was fixed across samples, so trajectories reflect redistribution of transcript abundance. **(b)** Effective expression breadth, calculated as the exponential of Shannon entropy; corresponding *q* values were < 0.001, < 0.001 and 0.02. Stars follow the thresholds defined in Fig. 4. These are RNA-allocation summaries, not direct measurements of translation or cellular energy state.

A focused S*. lycopersicum* case study provided a more detailed allocation decomposition (Online Resource 1: Fig. S12; Online Resource 2: Tables S58–S61). From 16–23 to 40–47 HPI, abundance-weighted CAI declined from 0.74 to 0.70, effective breadth increased from 907 to 1,274 genes, and the top-100-gene share declined from 50.4% to 38.6%. Abundance assigned to the lowest fixed-CAI quartile increased from 3.41% to 4.81%, while the highest-quartile share declined from 85.66% to 78.59%. Secreted/extracellular genes increased from 2.20% to 7.71% of relative transcript-molecule abundance, but their exclusion retained both the increase in low-residual-CAI abundance and the decline in weighted residual CAI (BH *q* < 0.01 for each). These patterns are consistent with changing transcript allocation; however, because we did not repeat this fuller decomposition in the other two datasets, it supports S. lycopersicum-specific inference.

A cross-system functional decomposition asked which non-translation genes contributed most strongly to each adjacent codon-demand change (Online Resource 1: Fig. S13; Online Resource 2: Tables S30, S33 and S35). After excluding 456 ribosome/translation-associated genes, 11,242 genes remained, and we evaluated the top 5% (563 per transition) against 4,999 abundance-, coding-length-, and change-stratified null sets. Across all 8,890 study-transition-term tests, 37 associations representing 16 GO terms passed global BH correction (*q* < 0.05) and occurred in nine of ten transitions. ATP-synthase activity or complex, ATP binding, and extracellular-region annotations recurred in all three host systems; TCA-cycle terms recurred in *S. lycopersicum* and *P. patens*, cellulose binding in *S. lycopersicum* and *A. thaliana*, and pectin catabolism in *A. thaliana* and *P. patens*. Exact terms and timing differed and GO terms overlap; these results identify functions among large RNA-level codon-demand contributors, not pathway activity or translation efficiency.

### Published infection programs connect global codon shifts to biological stage

In the *S. lycopersicum* source study, minor necrotic patches first appeared at 23 HPI and established necrotic growth was present at 40 HPI (Srivastava et al. 2020; Online Resource 2: Table S62). The largest *S. lycopersicum* codon-demand displacement and the greatest number of globally corrected functional associations occurred across this same 23–40-HPI interval. This temporal correspondence supplies macroscopic stage context but does not show that lesion development caused the codon-demand change. The source experiments supplied complementary biological context without directly measuring codon-dependent translation (Online Resource 1: Fig. S14d; Online Resource 2: Tables S62–S64). In *A. thaliana*, confocal imaging and fungal-biomass quantitative PCR separated attachment and germination, invasion and colonization, and later expansion and reproduction (Wei et al. 2024). Five of 48 tested fungal candidate proteins strongly induced cell death in *Nicotiana benthamiana*, but their mapped genes together represented less than 0.3% of fungal transcript-molecule allocation at every sampled time point. In *P. patens*, microscopy placed germination at 4 HPI, appressoria at 8 HPI, and invasive proliferation with tissue browning by 24 HPI; published RT-qPCR measurements were strongly concordant with RNA-seq for the selected pathogenesis transcripts at 8 and 24 HPI (Reboledo et al. 2021). These assays support developmental interpretation and selected RNA or candidate-protein observations. The clearest functional support came from transferring source-defined gene programs into independently generated target series. Wei modules M2, M3, and M6, defined in *A. thaliana*, contributed more to the *S. lycopersicum* first-to-last displacement than abundance-, coding-length-, and GC3-matched gene sets (28.7%, 15.4%, and 16.0% attenuation on removal; BH *q* < 0.01 each). M6 increased as *S. lycopersicum* lesions expanded and contains hydrolases, pectin esterases, and known cell-death-inducing proteins (Wei et al. 2024). Wei modules were also transferred to *P. patens*: M8 reduced 4–24-HPI displacement by 24.5% (*q* < 0.01), and M2 reduced the interpolated 16–24-HPI displacement by 29.0% (*q* < 0.05). Conversely, the Reboledo putatively secreted-protein differentially expressed gene set reduced the newer *A. thaliana* 16–24-HPI displacement by 13.4% (*q* < 0.01; Online Resource 1: Fig. S14c; Online Resource 2: Tables S65–S66).

Within-study decompositions were consistent with the same biological sequence. Early germination and appressorium modules and a late conidiation-associated module had disproportionate leverage in the *A. thaliana* trajectory; *P. patens* - and angiosperm-associated programs contributed to particular host series and intervals (Reboledo et al. 2021; Wei et al. 2024). By contrast, the broad candidate-effector and putatively secreted programs gained substantial transcript allocation but were not consistently exceptional in matched leave-set-out tests. Large allocation changes therefore need not produce unusual synonymous-demand displacement, and smaller coordinated programs can have disproportionate leverage. Full allocation trajectories, identifier reconciliation, selected-gene overlaps, and all study × set × contrast results are reported in Online Resource 2: Tables S48–S50 and S65–S66. The stricter source-set null confirmed the screening-significant findings without sparse-cell fallback. All 11 screening-significant cases remained significant after exact abundance-by-coding-length-by-GC3 matching with additional balance for expression-change magnitude and amino-acid composition; all displayed cases had strict BH *q* < 0.01 (Online Resource 2: Table S50). Because this better-balanced null identified additional cases rather than acting as a uniformly more conservative filter, inference remains tied to its stated matching conditions. Coherent signed induction or repression and amino-acid variation beyond the two balanced principal components can still distinguish a biological program from its controls.

### Host and targeted controls define the scope of inference

Synonymous-codon demand changed with time in each matched host (R² = 0.69, 0.67 and 0.91; all BH *q* < 0.01), and those effects survived gene-set and protein-family controls (Online Resource 1: Fig. S15; Online Resource 2: Table S67). None exceeded its host-specific matched trajectory null (empirical *P* = 0.35, 1.00 and 0.27; Fig. 6a; Online Resource 2: Table S68). Over the time-aligned 16–24-HPI contrast, host–fungus direction was weak and uncertain in *S. lycopersicum* (cosine = 0.10, bootstrap 95% CI −0.21–0.21) and *A. thaliana* (cosine = −0.23, bootstrap 95% CI −0.46–0.12), but positive in *P. patens* (cosine = 0.709, bootstrap 95% CI 0.69–0.72; Fig. 6b; Online Resource 1: Fig. S9; Online Resource 2: Tables S42 and S69). The first two intervals include zero; therefore, fungal demand does not seem to be a passive copy of a universal host codon-demand program.

**Figure 6.**
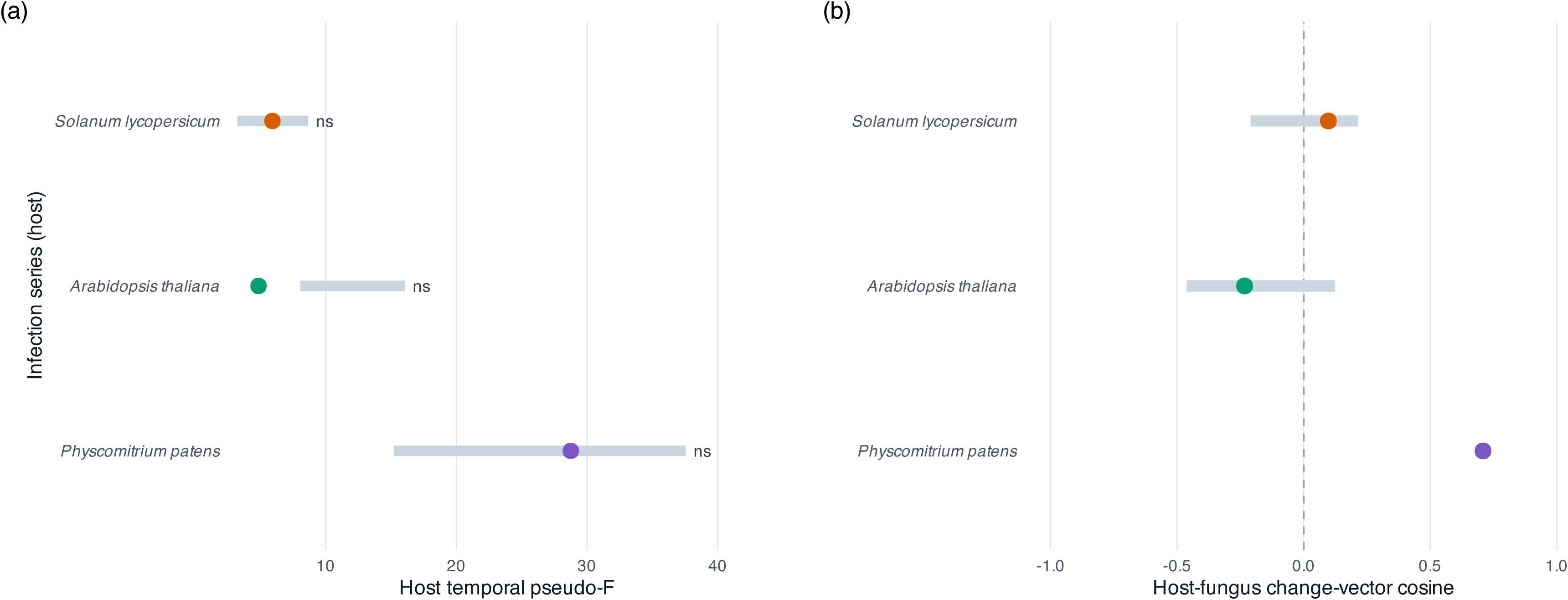
Matched hosts provide comparative context for fungal trajectories. **(a)** Observed host temporal pseudo-*F* values and 2.5th–97.5th percentiles from 1,999 host-specific matched trajectory reassignments. None of the three observed statistics exceeded their matched null after BH correction (*q*: *S. lycopersicum*, 0.53; *A. thaliana*, 1.00; *P. patens*, 0.53). **(b)** Host–fungus cosine similarity over the time-aligned 16–24-HPI interval: *S. lycopersicum*, 0.10 (95% bootstrap CI −0.21–0.21); *A. thaliana*, −0.23 (−0.46–0.12); and *P. patens*, 0.71 (0.69–0.72). Only the *P. patens* interval excludes zero. Host and fungal coding references were analyzed separately; the comparison assumes no shared codon preference or translation system and remains descriptive because host state, fungal state, and tissue composition covary.

The external sequence-calibration example was *BcNPTII*, a B05.10-codon-optimized neomycin phosphotransferase II selectable-marker coding sequence (Tang et al. 2024). The *BcNPTII* cassette occurs in the knockout vector; *BcBOT2* is the separate endogenous fungal gene targeted for deletion, not a gene contained within *BcNPTII*. Optimization moved the marker from low to high fungal CAI and demand percentiles (Online Resource 1: Fig. S16; Online Resource 2: Tables S70–S71). *BcBOT2* varied with time and served only as expression context (Online Resource 2: Table S72). Because the marker was designed using codon criteria and was not expressed in these infection datasets, this comparison shows that the metrics recognize and recover the potential effects of the redesigned sequence.

Alternative genomic tRNA-decoding rules retained strong temporal effects, although codon-capacity ranks were sensitive to the assumed wobble and inosine rules (Online Resource 1: Fig. S17; Online Resource 2: Tables S73–S75). The predefined all-gene GAA/GAG contrast showed greater 16–24-HPI organization than the function-conditioned null in S. lycopersicum and *P. patens* (BH *q* < 0.01 for each), but not in *A. thaliana* (*q* = 0.55); none of the three predefined contrasts passed correction within the secreted/extracellular candidate set (Online Resource 1: Fig. S18a; Online Resource 2: Table S76). Among 15 candidate tRNA-modification genes, time effects passed BH correction for 12 in *S. lycopersicum*, six in *A. thaliana,* and two in *P. patens* (Online Resource 1: Fig. S18b; Online Resource 2: Table S77). Local synonymous placement differed from within-gene randomization, especially among secreted/extracellular genes, yet only the *Arabidopsis* temporal contrast survived correction (Online Resource 1: Fig. S19; Online Resource 2: Tables S78–S80). These targeted results nominate GAA/GAG decoding and modification machinery for direct measurement, but transcript abundance is not enzyme activity, and none of these analyses measures tRNA modification, charging, or ribosome movement.

## Discussion

### Temporal transcriptome reallocation creates a dynamic codon-demand phenotype

Across all three infection time-series, progression through infection was accompanied by substantial reorganization of the synonymous codons represented in the *B. cinerea* transcript pool. Within each dataset, samples separated primarily by time in codon-demand space. The temporally aligned early transitions shared a positive directional component, whereas the longer trajectories diverged. Codon-preference-weighted abundance and expression breadth also changed over time, but not in a common direction or at the same time. Together, our results identify a recurrent transcript-level phenotype: as different genes become abundant, the aggregate synonymous composition of the message pool changes even though each gene’s coding sequence remains fixed. This perspective differs from the evolutionary framing usually applied to codon usage. Comparative studies describe genomic codon patterns as the joint outcome of nucleotide composition, drift, gene function, expression, and selection on decoding; in plant-pathogenic fungi, codon optimization has also been associated with the evolution of generalist parasitism (Badet et al. 2017; Baeza et al. 2024; LaBella 2024). Expression-weighted analyses instead ask which codons are represented in the active transcriptome at a particular state (Gingold et al. 2012). Our results therefore concern acclimation of transcript allocation during infection, not inherited codon adaptation or sequence evolution over hours.

A weighted codon profile will move whenever expression shifts among genes with different synonymous compositions, so temporal separation alone does not establish additional biological organization. The trajectory-reassignment tests addressed this issue by asking whether native expression trajectories were more strongly associated with their coding sequences than expected after matching genes for abundance and sequence properties. *S. lycopersicum* and *A. thaliana* exceeded that expectation across their complete series. *P. patens* did not across its full trajectory, but it did during the temporally aligned early interval. Temporal codon-demand change was therefore recurrent, whereas the additional organization attributable to native gene-to-sequence pairing depended on the infection system, interval, and null model. This distinction reconciles the shared early movement with the divergent later paths.

### A shared early component gives way to system-specific infection programs

The positive 16–24-HPI cosine similarities indicate that the early trajectories contained a common directional component, but HPI is only an approximate measure of fungal state. The source studies place germination, appressorium formation, penetration, tissue colonization, and lesion expansion at partly different times (Srivastava et al. 2020; Reboledo et al. 2021; Wei et al. 2024). Plant defense and lesion-development programs likewise overlap rather than proceed on a perfectly synchronized schedule (Windram et al. 2012). The early agreement therefore reflects recurrence across comparable infection windows, not proof that the fungi occupied identical developmental states or used the same mechanism at the same clock time.

Functional analyses connected these paths to the programs being deployed. Groups associated with germination and appressorium formation, fungal expansion, and later development contributed more strongly than expected to particular transitions. Programs defined in the Wei *A. thaliana* experiment also contributed in the independent *S. lycopersicum* and *P. patens* datasets, while the Reboledo putatively secreted set contributed to the newer *A. thaliana* series (Reboledo et al. 2021; Wei et al. 2024). Testing sets outside the experiments in which they were defined reduces circularity and suggests that related infection programs recur across datasets. Moreover, all initially significant associations survived the strict 102-test analysis, which matched focal and control genes for expression, coding length, GC3, expression-change magnitude, and amino-acid composition. Broad abundance or sequence differences between functional sets are therefore unlikely to explain these associations, although the evidence remains RNA-based.

The GO analysis provided a complementary view: ATP-synthase or respiratory functions and extracellular annotations recurred across hosts, whereas pectin, chromatin, lipid, and amino-acid functions appeared in particular intervals. This mixture of shared core processes and system-specific modules echoes a broader cross-host analysis of *B. cinerea* (Singh et al. 2026), while fungal conversion of plant hexoses into mannitol during sunflower infection illustrates changing carbon use during disease development (Dulermo et al. 2009). Neither parallel identifies a single causal virulence mechanism. Secreted phytotoxic factors can be highly redundant (Leisen et al. 2022), and overlapping GO enrichments establish neither pathway flux nor protein synthesis, secretion, or necessity. The primary *S. lycopersicum* series illustrates this stage-based interpretation, where its largest transition separated minor necrotic patches at 23 HPI from established necrotic growth at 40 HPI. However, the 23-and 47-HPI samples were collected under red light before lights-on (Srivastava et al. 2020), and light affects *B. cinerea* development and virulence (Canessa et al. 2013). Developmental progression, illumination, and diel phase therefore cannot be separated.

### A resource-allocation hypothesis

Later *S. lycopersicum* and *A. thaliana* samples combined a broader effective gene repertoire with lower mean CAI calculated against ribosomal reference genes. This pattern is consistent with a shift from concentrated investment in a narrower transcript set toward a more diverse infection program. *P. patens* instead reached a transient codon-preference maximum before breadth increased, arguing against a universal progression from more-preferred to less-preferred codons. The relationship therefore depends on the program and stage sampled, and lower ribosomal-reference CAI alone demonstrates neither slower translation nor greater energetic cost.

Resource allocation nevertheless provides a testable hypothesis. Nitrogen availability alters *B. cinerea* infection and virulence-associated expression (Soulié et al. 2020), while yeast studies show condition-dependent redistribution of proteome and translational capacity (Metzl-Raz et al. 2017; Yu et al. 2020). Codon optimality influences mRNA decay in budding yeast (Presnyak et al. 2015), and codon usage can affect transcription in *Neurospora* (Zhou et al. 2016). Changes in tRNA abundance or modification can redirect protein synthesis, protect proteome integrity, and regulate effector secretion or ergosterol-linked virulence in *Magnaporthe oryzae* (Nedialkova and Leidel 2015; Torrent et al. 2018; Li et al. 2023; He et al. 2024). Therefore, although many mechanistic precedents exist, no evidence shows that the same mechanisms produced our trajectories. Future work in *B. cinerea* should consider whether variation in resource allocation corresponds to virulence phenotypes, and whether allometric investment in the tRNA pool is present.

In our data, GAA/GAG organization in *S. lycopersicum* and *P. patens*, and stage-dependent expression of candidate tRNA-modification genes, suggest links to decoding biology. Their absence from the *A. thaliana* contrast and secreted-gene subset argues against a single conserved response. No protein, ribosome, tRNA, nutrient, or ATP measurements were available to distinguish transcript allocation from RNA-stability or translational effects. Our analysis placed the deliberately recoded *BcNPTII* marker in the high-preference region (Tang et al. 2024), and previous *B. cinerea* reporters show that synonymous design can alter RNA abundance and fluorescence (Leroch et al. 2011).

### Robustness and scope of cross-study inference

Technical sensitivity analyses make read assignment an unlikely explanation for the temporal structure. The principal trajectories survived effective-length normalization, unique-only competitive mapping, host-homology exclusion, and mock-derived gene blacklists. The elevated signal in the *P. patens* 8-h mock remained unexplained, but its genes scarcely overlapped the host-homology mask, and removing every gene with mock signal did not eliminate the *P. patens* trajectory. Conversely, the primary *S. lycopersicum* study lacks a matched pure-host control, leaving an assignment limitation that computation cannot remove.

Sequence concordance further narrows the claim. Expressed coding positions in *S. lycopersicum* and *A. thaliana* were nearly identical to B05.10; the lemon-derived *P. patens* isolate was highly concordant but detectably different, and incorporating its RNA-supported substitutions did not change the result. We therefore show recurrence among *B. cinerea* infection datasets represented in B05.10 coordinates, not three authenticated replicates of one genotype. RNA-seq variants cover only sufficiently expressed positions and cannot replace whole-genome genotyping (Piskol et al. 2013). Host trajectories also failed to exceed matched reassignments, while host–fungus directions ranged from weakly opposed to aligned. Fungal movement was not a passive copy of one universal whole-sample direction, and separate translation systems preclude inference of direct host–fungus competition for tRNAs.

Biological interpretation remains more limited than detection of the temporal pattern. Gene-level length correction approximates relative transcript molecules but lacks isoform inference and full sequence, positional, and fragment-GC bias modeling (Patro et al. 2017). RNA abundance cannot separate transcription from degradation or identify ribosome-engaged messages. Host, isolate, tissue, protocol, and study are partly confounded. These constraints preclude a replicated host effect, a universal direction, or a mechanistic conclusion about translation, but not the narrower finding that infection-associated transcript reallocation repeatedly changes fungal synonymous-codon representation.

### From dynamic demand to causal tests

Future work needs to measure these links in the same infection material, whether in organs, hosts, or simulated infections on media plates. External RNA standards with fungal biomass estimates would distinguish absolute from compositional allocation (Lovén et al. 2012), and imaging would align samples by fungal state. Ribosome profiling and proteomics would test translation (Ingolia et al. 2009), while quantitative tRNA profiling would measure abundance, charging, and modification (Behrens et al. 2021). For example, a complementary reporter could encode one protein with early-demand-matched or late-demand-matched synonymous sequences while holding promoter, untranslated regions, copy number, and integration site constant. RNA half-life, ribosome occupancy, protein accumulation, and infection performance would then separate stability from translational effects.

This framework adds sequence composition to conventional functional genomics: pathway analysis identifies which programs change, whereas transcriptome-weighted codon demand describes a consequence of deploying them. Applying both approaches to a panel of isolate-specific time-series transcriptomic datasets may separate within-isolate acclimation from inherited coding differences (Atwell et al. 2015; Soltis et al. 2020). Here, synonymous codon demand emerged as a dynamic property of the inferred *B. cinerea* transcript pool whose reorganization recurred across infection systems but followed program- and interval-specific paths. That position between differential expression and direct translation measurements generates testable hypotheses about RNA stability, decoding, and resource allocation without treating those mechanisms as demonstrated.

## Statements and declarations

## Author contributions

JAD designed the study and collected and analyzed the data. JAD wrote the manuscript with input from LP and MM.

## Data availability

Raw RNA-seq reads are publicly available under BioProject accessions PRJNA628162, PRJNA1158549, PRJNA647932 and PRJNA1056687. The B05.10 reference is available under assembly accession GCF_000143535.2; host references and exact versions are listed in Online Resource 1, Supplementary Methods SM2. The *BcNPTII* sequence is available under GenBank accession OR921207.

## Code availability

Analysis code, processed matrices, manifests, seeds, software environments, and figure-generating inputs will be archived at **[add GitHub]**.

## Supplementary information

Online Resource 1 (ESM_1.pdf) contains Supplementary Methods SM1–SM10 and Figs. S1– S19 with legends. Online Resource 2 (ESM_2.xlsx) contains Tables S1–S80 and the machine-readable manifest.

## Supporting information

Supplmental Resource 1

Supplmental Resource 2

## References

1. Aitchison J (1986) The statistical analysis of compositional data. Chapman & Hall, London

2. Alonge M, Lebeigle L, Kirsche M et al (2022) Automated assembly scaffolding using RagTag elevates a new S. lycopersicum system for high-throughput genome editing. Genome Biol 23:258. 10.1186/s13059-022-02823-7

3. Anderson MJ (2001) A new method for non-parametric multivariate analysis of variance. Austral Ecol 26:32–46. 10.1111/j.1442-9993.2001.01070.pp.x

4. Atwell S, Corwin J, Soltis N, Subedy A, Denby K, Kliebenstein DJ (2015) Whole-genome resequencing of *Botrytis cinerea* isolates identifies high levels of standing diversity. Front Microbiol 6:996. 10.3389/fmicb.2015.00996

5. Badet T, Peyraud R, Mbengue M et al (2017) Codon optimization underpins generalist parasitism in fungi. eLife 6:e22472. 10.7554/eLife.22472

6. Baeza M, Sepulveda D, Cifuentes V, Alcaíno J (2024) Codon usage bias in yeasts and its correlation with gene expression, growth temperature and protein structure. Front Microbiol 15:1414422. 10.3389/fmicb.2024.1414422

7. Behrens A, Rodschinka G, Nedialkova DD (2021) High-resolution quantitative profiling of tRNA abundance and modification status in eukaryotes by mim-tRNAseq. Mol Cell 81:1802–1815.e7. 10.1016/j.molcel.2021.01.028

8. Canessa P, Schumacher J, Hevia MA, Tudzynski P, Larrondo LF (2013) Assessing the effects of light on differentiation and virulence of the plant pathogen *Botrytis cinerea*: characterization of the White Collar Complex. PLoS One 8:e84223. 10.1371/journal.pone.0084223

9. Cheung N, Tian L, Liu X, Li X (2020) The destructive fungal pathogen *Botrytis cinerea*— insights from genes studied with mutant analysis. Pathogens 9:923. 10.3390/pathogens9110923

10. Dahal U, Shakya R, Kour B, Khanal B, Singh B (2026) Evolution and functional implications of codon usage bias in eukaryotes. J Mol Evol 94:286–303. 10.1007/s00239-026-10305-8

11. Dulermo T, Rascle C, Chinnici G et al (2009) Dynamic carbon transfer during pathogenesis of sunflower by the necrotrophic fungus *Botrytis cinerea*: from plant hexoses to mannitol. New Phytol 183:1149–1162. 10.1111/j.1469-8137.2009.02890.x

12. Gingold H, Dahan O, Pilpel Y (2012) Dynamic changes in translational efficiency are deduced from codon usage of the transcriptome. Nucleic Acids Res 40:10053–10063. 10.1093/nar/gks772

13. Goormaghtigh F, and Bumann D (2026) Stress-adapted codon usage enables efficient virulence-gene expression in Salmonella. bioRxiv 2026.02.22.707322. 10.64898/2026.02.22.707322

14. Gupta S, Paul K, Roy A (2021) Codon usage signatures in the genus *Cryptococcus*: a complex interplay of gene expression, translational selection and compositional bias. Genomics 113:821–830. 10.1016/j.ygeno.2020.10.013

15. He RR, Lv Z, Li Y et al (2024) tRNA-m1A methylation controls the infection of *Magnaporthe oryzae* by supporting ergosterol biosynthesis. Dev Cell 59:2931–2946.e7. 10.1016/j.devcel.2024.08.002

16. Ingolia NT, Ghaemmaghami S, Newman JRS, Weissman JS (2009) Genome-wide analysis in vivo of translation with nucleotide resolution using ribosome profiling. Science 324:218–223. 10.1126/science.1168978

17. LaBella AL (2024) A symphony of roles for codon usage in fungal genomics. In: Nowrousian M, Stajich JE (eds) Fungal genomics. The Mycota, vol 13. Springer, Cham, pp 75–100. 10.1007/978-3-031-75666-5_6

18. Leisen T, Werner J, Pattar P et al (2022) Multiple knockout mutants reveal a high redundancy of phytotoxic compounds contributing to necrotrophic pathogenesis of *Botrytis cinerea*. PLoS Pathog 18:e1010367. 10.1371/journal.ppat.1010367

19. Leroch M, Mernke D, Koppenhoefer D et al (2011) Living colors in the gray mold pathogen *Botrytis cinerea*: codon-optimized genes encoding green fluorescent protein and mCherry, which exhibit bright fluorescence. Appl Environ Microbiol 77:2887–2897. 10.1128/AEM.02644-10

20. Li G, Dulal N, Gong Z, Wilson RA (2023) Unconventional secretion of *Magnaporthe oryzae* effectors in rice cells is regulated by tRNA modification and codon usage control. Nat Microbiol 8:1706–1716. 10.1038/s41564-023-01443-6

21. Lovén J, Orlando DA, Sigova AA et al (2012) Revisiting global gene expression analysis. Cell 151:476–482. 10.1016/j.cell.2012.10.012

22. Metzl-Raz E, Kafri M, Yaakov G, Soifer I, Gurvich Y, Barkai N (2017) Principles of cellular resource allocation revealed by condition-dependent proteome profiling. eLife 6:e28034. 10.7554/eLife.28034

23. Nedialkova DD, Leidel SA (2015) Optimization of codon translation rates via tRNA modifications maintains proteome integrity. Cell 161:1606–1618. 10.1016/j.cell.2015.05.022

24. Patro R, Duggal G, Love MI, Irizarry RA, Kingsford C (2017) Salmon provides fast and bias-aware quantification of transcript expression. Nat Methods 14:417–419. 10.1038/nmeth.4197

25. Piskol R, Ramaswami G, Li JB (2013) Reliable identification of genomic variants from RNA-seq data. Am J Hum Genet 93:641–651. 10.1016/j.ajhg.2013.08.008

26. Presnyak V, Alhusaini N, Chen YH et al (2015) Codon optimality is a major determinant of mRNA stability. Cell 160:1111–1124. 10.1016/j.cell.2015.02.029

27. Reboledo G, Agorio A, Vignale L, Batista-García RA, Ponce de León I (2021) *Botrytis cinerea* transcriptome during the infection process of the bryophyte *Physcomitrium patens* and angiosperms. J Fungi 7:11. 10.3390/jof7010011

28. Sharp PM, Li WH (1987) The Codon Adaptation Index—a measure of directional synonymous codon usage bias, and its potential applications. Nucleic Acids Res 15:1281–1295. 10.1093/nar/15.3.1281

29. Shi H, Ding G, Wang Y et al (2025) Genome-wide identification of long non-coding RNA for *Botrytis cinerea* during infection to S. lycopersicum (*Solanum lycopersicum*) leaves. BMC Genomics 26:7. 10.1186/s12864-024-11171-8

30. Singh R, Muhich AJ, Tom C et al (2026) Combined generalist and host-specific transcriptional strategies enable host generalism in the fungal pathogen *Botrytis cinerea*. Proc Natl Acad Sci USA 123:e2521414123. 10.1073/pnas.2521414123

31. Soltis NE, Caseys C, Zhang W, Corwin JA, Atwell S, Kliebenstein DJ (2020) Pathogen genetic control of transcriptome variation in the *Arabidopsis thaliana*–*Botrytis cinerea* pathosystem. Genetics 215:253–266. 10.1534/genetics.120.303070

32. Soulié MC, Koka SM, Floch K et al (2020) Plant nitrogen supply affects the *Botrytis cinerea* infection process and modulates known and novel virulence factors. Mol Plant Pathol 21:1436–1450. 10.1111/mpp.12984

33. Srivastava DA, Arya GC, Pandaranayaka EPJ et al (2020) Transcriptome profiling data of *Botrytis cinerea* infection on whole plant *Solanum lycopersicum*. Mol Plant Microbe Interact 33:1103–1107. 10.1094/MPMI-05-20-0109-A

34. Tang M, Wang Y, Wang K et al (2024) Codon optimization enables the geneticin resistance gene to be applied efficiently to the genetic manipulation of the plant pathogenic fungus *Botrytis cinerea*. Plants 13:324. 10.3390/plants13020324

35. Torrent M, Chalancon G, de Groot NS, Wuster A, Babu MM (2018) Cells alter their tRNA abundance to selectively regulate protein synthesis during stress conditions. Sci Signal 11:eaat6409. 10.1126/scisignal.aat6409

36. Van Kan JAL, Stassen JHM, Mosbach A et al (2017) A gapless genome sequence of the fungus *Botrytis cinerea*. Mol Plant Pathol 18:75–89. 10.1111/mpp.12384

37. Wei J, Zhou Q, Zhang J, Wu M, Li G, Yang L (2024) Dual RNA-seq reveals distinct families of co-regulated and structurally conserved effectors in *Botrytis cinerea* infection of *Arabidopsis thaliana*. BMC Biol 22:239. 10.1186/s12915-024-02043-4

38. Windram O, Madhou P, McHattie S et al (2012) *Arabidopsis* defense against *Botrytis cinerea*: chronology and regulation deciphered by high-resolution temporal transcriptomic analysis. Plant Cell 24:3530–3557. 10.1105/tpc.112.102046

39. Wright F (1990) The effective number of codons used in a gene. Gene 87:23–29. 10.1016/0378-1119(90)90491-9

40. Yu R, Campbell K, Pereira R et al (2020) Nitrogen limitation reveals large reserves in metabolic and translational capacities of yeast. Nat Commun 11:1881. 10.1038/s41467-020-15749-0

41. Zhou Z, Dang Y, Zhou M et al (2016) Codon usage is an important determinant of gene expression levels largely through its effects on transcription. Proc Natl Acad Sci USA 113:E6117–E6125. 10.1073/pnas.1606724113

42. Zhou Z, Dang Y, Zhou M, et al (2018) Codon usage biases co-evolve with transcription termination machinery to suppress premature cleavage and polyadenylation. eLife. 7:e33569. 10.7554/eLife.33569

43. Zhou Z, Dang Y, Zhou M, Yuan H, Liu Y. 2018. Codon usage biases co-evolve with transcription termination machinery to suppress premature cleavage and polyadenylation. Jensen TH, editor. eLife. 7:e33569. doi:10.7554/eLife.33569.

