## Supplementary material for "Transcriptome reallocation during infection reshapes synonymous codon demand in *Botrytis cinerea*": Supplmental Resource 1

Supplementary Methods SM1-SM10 and Figures S1-S19

##### Supplementary figure inventory


| | |
|---|---|
| **Figure S1.** Robustness to protein-family weighting and translation-gene exclusion | **Figure S2.** Construction of the time-aligned 16–24-HPI contrast |
| **Figure S3.** Matched expression-trajectory reassignment null | **Figure S4.** Shared early-window codon-demand organization and displacement |
| **Figure S5.** Functional-group-conditioned trajectory reassignment | **Figure S6.** Effective-length sensitivity of temporal codon demand |
| **Figure S7.** Temporal inference is tested under strict cross-mapping and mock-host controls | **Figure S8.** Expressed coding sites quantify reference concordance without authenticating isolate identity |
| **Figure S9.** Bootstrap uncertainty in time-aligned directional cosines | **Figure S10.** *Solanum lycopersicum* (supplementary) sensitivity analysis |
| **Figure S11.** Independent ribosomal-reference CAI analysis | **Figure S12.** Tomato-series detailed transcript-allocation metrics |
| **Figure S13.** Non-translation programs reshape synonymous codon demand | **Figure S14.** Source-study experiments and gene programs contextualize codon-demand reorganization |
| **Figure S15.** Matched host robustness and trajectory nulls | **Figure S16.** BcNPTII external sequence-calibration example |
| **Figure S17.** Sensitivity to genomic tRNA decoding assumptions | **Figure S18.** Predefined modification-sensitive codon and machinery tests |
| **Figure S19.** Local synonymous placement near coding-sequence starts | |

### Supplementary methods

#### SM1. Public datasets, sample inclusion and study roles

Four public dual-RNA-seq datasets were evaluated against one common B05.10 coding reference. Three well-replicated datasets were designated as independent primary systems: PRJNA628162 in *Solanum lycopersicum* at 16, 23, 40 and 47 hours post inoculation (HPI); PRJNA1158549 in *Arabidopsis thaliana* at 6, 12, 24, 48, 72 and 96 HPI; and PRJNA647932 in *Physcomitrium patens* at 4, 8 and 24 HPI. Each primary time point contained three biological libraries. The tomato and *Arabidopsis* source studies explicitly name B05.10. The moss study describes a *B. cinerea* strain isolated from lemon and maps it to the B05.10 reference, but does not explicitly identify the inoculum as B05.10. PRJNA1056687 contained two tomato libraries at 6, 12, 24, and 48 HPI and was retained as a supplementary sensitivity dataset rather than combined with the primary evidence. Reference consistency was assessed with an expressed-CDS RNA-seq variant panel as described in SM2; this is a concordance sensitivity rather than whole-genome isolate authentication.

Only infection libraries from the selected experiments were analyzed. We excluded uninfected host, fungal culture, and other treatment libraries. The three deposited 0-HPI libraries associated with PRJNA628162 are synthetic mixtures of mock-inoculated whole-leaf RNA and B05.10 spore RNA; they were excluded from all infection-time tests and retained only in archived sensitivity outputs. Table S1 records run accession, host genotype or accession, sampled material, time, replicate, library layout, biological-unit definition, and inclusion status for all 47 analyzed infection libraries and the three excluded mixtures. Cross-study design and primary temporal statistics are reported in Tables S2 and S40–S41, while the lower-replication tomato sensitivity is reported in Tables S3 and S54 and Figure S10.

#### SM2. Competitive host–fungus quantification

The B05.10 fungal reference was derived from GCF_000143535.2. For PRJNA628162, single-end reads were mapped with Rsubread `subjunc` to a species-prefixed combined M82 GCA_027704885.1 plus B05.10 GCF_000143535.2 genomic reference. Phred+33 scores were used; mapping logs record three allowed mismatches, a maximum of 5-nt indels, multimapping detection, and at most one reported alignment. Fungal exon counts were summarized unstranded by gene with `featureCounts`, excluding multimapping and multi-overlap assignments. Primary tomato host abundance was generated separately with Rsubread `align` against source-provided high-confidence representative M82 cDNAs plus B05.10 CDS competitors and counted unstranded at mapping quality ≥1.

PRJNA1056687, PRJNA1158549 and PRJNA647932 used paired-end competitive alignments to host RNA plus B05.10 transcript references. `featureCounts` required both read ends to map, used mapping quality ≥1, and excluded multimapping and multi-overlap assignments; a proper-pair flag was not explicitly required. The fungal transcript reference retained the longest CDS per locus tag. The PRJNA1056687 libraries came from cv. Moneymaker but were mapped against the available M82 representative transcript reference, so its host-side results carry an explicit cultivar/reference mismatch and remain supplementary. M82 supplied one representative transcript and CDS per high-confidence locus. Arabidopsis and moss mapping used their corresponding RefSeq RNA records grouped by parsed gene identifier, while downstream host codon profiles retained the longest valid CDS per gene. Thus, reads were assigned competitively within each mapping workflow, but the primary tomato fungal and host matrices came from separate genomic and transcriptomic alignments.

PRJNA628162 and PRJNA1056687 logs record Rsubread 2.22.1; the saved primary-tomato mapping session records R 4.5.2. Every PRJNA1158549 and PRJNA647932 alignment log directly records Rsubread 2.26.0. Their mapping R version was retrospectively reconstructed as R 4.6.0 with high confidence

For PRJNA628162, competitive fungal counts were compared with the original fungus-only workflow. The comparison included assigned counts, per-library count correlations, species-level alignment records, and mapping-quality summaries (Tables S4–S7). We competitively mapped the external primary systems from the outset. Study-specific host assignment quality is summarized in Table S5.

A stricter cross-assignment sensitivity realigned infection libraries to one-host-plus-B05.10 transcript references with unique alignments only and counted at MAPQ thresholds 1, 20 and 40. The fungal component contained one longest CDS per gene, preventing alternative fungal isoforms from being treated as cross-species ambiguity. Three uninfected Arabidopsis libraries and one mock-inoculated moss library at each of 4, 8 and 24 HPI were mapped identically to estimate false fungal assignment and define conservative fungal-gene blacklists; the primary tomato study lacked a pure host-only control because its 0-HPI deposits are host-plus-spore mixtures. Separately, local nucleotide searches identified B05.10 genes with host matches at ≥90% identity across ≥50 nt, with 85%/75-nt and 80%/100-nt alternatives. Whole genes meeting these criteria were excluded in conservative reruns. Figure S7 and Tables S13–S17 and S52 report the results.

We assessed reference consistency and isolate comparability using RNA-seq reads from three predeclared high-fungal-depth libraries per primary study. Up to 10 million read pairs or reads per library were uniquely mapped to the corresponding host-plus-B05.10 transcript reference with at most six mismatches. Candidate coding sites excluded the first and last ten bases of each CDS and required MAPQ ≥20, base quality ≥30, pooled depth ≥30, a major-allele fraction ≥0.90 and support from at least two libraries with per-library depth ≥5 and major-allele fraction ≥0.80. Reference concordance was summarized within study, between study consensuses, and across sites callable in all systems. The analysis covers expressed coding positions, is subject to reference-mapping bias, and cannot authenticate unexpressed or noncoding regions (Figure S8; Tables S18–S23).

For a direct inventory sensitivity, we jointly applied all 1,043 supported moss consensus substitutions to the corresponding B05.10 coding sequences after verifying every reference base and complete CDS. Codons containing more than one substitution would have been edited jointly; none occurred. All corrected sequences retained their reading frame and lacked internal stops, and we did not skip any supported site. Corrected per-gene codon counts replaced the B05.10 counts for the affected genes, while non-callable positions remained reference bases. The nine moss libraries were then reanalyzed with the same coding-length correction, within-family CLR response, and one fixed set of 9,999 label permutations for both inventories. Sample-level CLR profiles, distance matrices and first-to-last change vectors were compared directly (Table S24). This partial correction is a sensitivity to observed expressed-CDS differences, not a reconstruction of the isolate genome.

#### SM3. Coding-sequence metrics and codon-demand construction

Coding sequences were required to have valid reading frames and compatible gene identifiers. Sense codons were counted per gene. Wright effective number of codons (ENC) described gene-level codon-use concentration, and GC3 described synonymous third-position composition. Ribosomal-reference CAI weights were estimated once from 87 annotated B05.10 ribosomal genes and were fixed across libraries and studies. The fixed-score CAI analysis and annotation coverage are detailed in Tables S25 and S31 and Figure S11.

A composition-preserving null randomized synonymous placement while holding each protein sequence and genome-wide codon totals fixed. It evaluated whether ENC, GC3-adjusted ENC variation, and the CAI–expression association exceeded expectations from amino-acid composition and overall codon availability (Table S26).

Raw gene counts were divided by the analyzed coding-sequence length in codons and then closed to a constant library sum. This TPM-like quantity is a proxy for relative transcript-molecule abundance; using gene-level counts per million directly would count coding length once through fragment yield and again through the gene's codon inventory. We calculated library-specific fungal codon demand by multiplying relative transcript-molecule abundance by each gene's codon-count vector and summing across genes. Within each amino-acid family, demand was divided by total demand for that amino acid. A pseudocount of 1 × 10^-10 was added before centered log-ratio transformation. The primary response contained 59 synonymous-codon coordinates and excluded changes among amino acids. Amino-acid and synonymous summaries were retained separately throughout the workflow.

The length-denominator sensitivity held competitive gene counts fixed and compared three definitions on a common eligible gene set: analyzed CDS length, the exact countable annotation feature length supplied to `featureCounts`, and effective feature length calculated as feature length minus the library-specific mean fragment or read length plus one. Paired-end mean fragment lengths were recovered from the original Rsubread logs; the single-end PRJNA628162 read length was exactly 84 nt, calculated from SRA total bases divided by total spots. This sensitivity corrects feature-end opportunity but does not perform transcript-isoform expectation maximization or sequence-, position- or GC-bias correction. Figure S6 and Tables S8–S11 report temporal tests, profile concordance, retained genes and library-specific parameters.

#### SM4. Temporal tests and robustness analysis

Within each study, a permutational multivariate analysis tested time-point separation in the 59-coordinate fungal CLR profiles. The pseudo-$F$ statistic compared between-time-point with within-time-point variation, and $R^2$ described the fraction of variation associated with sampling time. Library labels were permuted 9,999 times, with each deposited biological library as the experimental unit. A permutation test of distances to time-point centroids assessed dispersion heterogeneity. We analyzed individual codons using ordinary least-squares models with time point as a categorical predictor. Overall temporal effects used the model $F$ test, and adjacent-time-point comparisons used $t$ contrasts based on the model's pooled residual variance. Benjamini–Hochberg adjustment was applied separately within study across the 59 overall codon tests and across every adjacent codon-by-transition contrast. Dispersion, adjacent-codon, early–late and family-balanced results are provided in Tables S34 and S36–S39; Figure S9 gives bootstrap uncertainty for the cross-dataset directional comparisons.

Robustness scenarios required fungal expression above alternative thresholds, excluded short coding sequences, weighted genes by inverse protein-family size at 50%, 70% and 90% identity, excluded the ribosomal reference, or excluded all annotated translation-associated genes. Each original library was omitted in turn. Counts were also downsampled 100 times to the lowest fungal depth within each study. Table S27 and its subset in Table S32 report an independent 9,999-permutation robustness-stage run; its baseline pseudo-$F$ and $R^2$ are identical to the canonical analysis in Table S2, while small differences in empirical $P$ reflect independent finite Monte Carlo draws. Current cross-study robustness, depth, and omission results are contained in Tables S2, S27, S32, and S40–S41 and Figure S1. Functional driver analyses are reported separately in SM6.

We tested individual synonymous codons using a categorical timepoint model. Overall probabilities were adjusted across 59 codons. Adjacent-timepoint contrasts used the pooled residual variance of the corresponding codon model and were adjusted across every codon-by-transition contrast in that study. Table S55 provides complete overall tests for all three primary systems; Figure 3 displays the descriptive 20-codon subset. Superseded tomato-only heatmaps and controlled CAI–abundance slopes are retained in the analysis archive rather than the consolidated submission supplement.

#### SM5. Matched trajectory reassignment and shared-window statistics

Matched trajectory reassignment tested whether changing expression automatically generated the observed codon-demand organization. Complete library trajectories were exchanged among genes within strata defined by mean expression, coding length, GC3, and the first two amino-acid-composition principal components. Thus, every null dataset retained the observed coding sequences and expression trajectories but broke their gene-specific pairing. The primary full-series null is provided in Figure S3 and Table S37.

The time-aligned early comparison used a 16–24-HPI linear contrast. No dataset observed both endpoints directly. We applied interpolation weights only to observed time-point means: 16, 23, and 40 HPI for tomato; 12 and 24 HPI for *A. thaliana*; and 8 and 24 HPI for *P. patens*. Interpolated values were not treated as new replicates or evidence that developmental state was equivalent. Euclidean displacement was the norm of the resulting change vector. Organization Q was defined as squared displacement divided by the sum of coordinate-wise replicate-estimated contrast variances (a trace-standardized displacement, not a Mahalanobis distance). Figure S2 visualizes the observed sampling grids and signed interpolation weights; Figure S4 and Tables S43–S45 report null distributions, weights, and contributing libraries. Directional cosine uncertainty was estimated with 10,000 resamplings of biological libraries within each observed time point (Figure S9; Table S42).

A stricter reassignment additionally required the donor and recipient genes to share a broad functional class: translation/RNA decoding, secreted/extracellular, metabolism, other annotated, or unannotated. Figure S5 and Tables S46–S47 report full-series and shared-window statistics and demonstrate the fraction of genes remaining movable within the conditioned strata.

#### SM6. Expression allocation and functional analyses

We summarized each fungal library by relative-transcript-molecule-abundance-weighted ribosomal-reference CAI, effective expression breadth, abundance shares assigned to fixed CAI quartiles, and the top-100-gene share. Global time effects were tested by label permutation. Early–late tomato contrasts used the exact assignment distribution and false-discovery correction across allocation metrics. Figure S12 and Tables S56–S59 contain the cross-study and detailed tomato allocation results.

Functional contributions were examined separately for every adjacent transition in the three primary infection series after removing 456 genes annotated for ribosome- or translation-related functions. For each transition, we decomposed the change in within-amino-acid synonymous codon shares into gene-level vectors and selected the top 5% of the 11,242 retained genes (563 genes) by absolute projection onto the observed change vector. GO terms among these large contributors were evaluated against 4,999 random gene sets drawn within joint 5 × 5 × 4 rank strata of study-wide mean relative transcript-molecule abundance, analyzed CDS length, and transition-specific absolute log2 abundance change. This is coarsened stratified matching rather than exact continuous matching; continuous-scale balance diagnostics are retained with the analysis. Benjamini–Hochberg adjustment was calculated within transition, within study, and across all 8,890 study-transition-term tests. Annotation coverage, standard over-representation results, and the matched analysis are reported in Figure S13 and Tables S30, S33, and S35. Secreted/extracellular genes were evaluated separately to determine whether their changing expression share accounted for the late lower-CAI tomato pattern (Tables S60–S61). Superseded count-weighted exploratory summaries and corrected adjacent-timepoint CAI–abundance slope contrasts are retained outside the submission package in the analysis archive and are not used in the current inferential chain.

We imported source-defined fungal programs from Wei et al. (nine Arabidopsis infection modules, 591 candidate effector proteins, the complete experimentally screened candidate set, and the five uniquely mapped candidates reported as strong cell-death inducers) and Reboledo et al. (host-shared, moss-associated, angiosperm-associated, and putatively secreted programs). The Reboledo moss/angiosperm categories were defined by differential expression against PDA-grown fungus at source-selected windows (moss 8/24 HPI, Arabidopsis 12/24 HPI, tomato 16/23 HPI and lettuce 48 HPI); they are source-classified expression programs, not causal tests of host specificity. Source identifiers were reconciled to the current B05.10 coding reference; membership and reconciliation coverage are audited in Tables S48–S49. Allocation of relative transcript-molecule abundance to each set was calculated in every library (Table S65). For the first-to-last observed contrast and the interpolated 16–24-HPI contrast, the full pooled-timepoint CLR displacement was compared with displacement after removing each source set. In the screening analysis, we evaluated each observed attenuation against 999 gene sets selected from 4 × 4 × 4 coarsened cells of mean abundance, CDS length, and GC3. The algorithm prioritized the exact target cell but used nearest cells, and then any eligible nontarget gene, if exact-cell capacity was insufficient; sets with fewer than ten mapped genes were summarized descriptively but not assigned a matched-set probability (Table S66). Benjamini–Hochberg adjustment was applied separately within each contrast and source-paper family (Wei or Reboledo) across target studies and gene sets. A stricter sensitivity preserved the target's exact 64-cell membership in every draw and additionally required balance for the three continuous base variables, contrast-specific absolute change on a log1p relative-abundance scale and amino-acid-composition PC1–PC2. We used randomized nearest initialization followed by legal within-cell swaps and accepted a draw only when the maximum absolute standardized mean difference was ≤0.10 and the root-mean-square standardized mean difference was ≤0.05. The strict analysis covered the complete original 102-test universe rather than only displayed cases, retained the original four multiplicity families, and recorded infeasible cases instead of relaxing the matching rules (Table S50). The added change covariate matched magnitude, not signed induction or repression, and did not constrain coordination among genes within a set. Because these matching schemes condition on selected summaries rather than every possible sequence or expression feature, significant attenuation denotes disproportionate contribution under the stated null rather than uniquely codon-compositional leverage. Reboledo applications to the current moss and primary tomato series partly reuse source RNA-seq; transfer to the newer Wei Arabidopsis series is independent at the RNA-seq-study level but not at the host-species or assay level. These analyses are functional decompositions, not independent translation validation.

Assay-level evidence and source-design information reported by the source studies were cataloged separately (Table S62). Published RT-qPCR and RNA-seq log2 fold changes for 16 moss pathogenesis genes were transcribed from Reboledo et al. Supplementary Table S5; both assays used PDA-grown fungus as the fold-change reference, and each reported value summarized three biological replicates. The source paper reported regression $R^2 = 0.9882$, whereas the present descriptive reanalysis calculated Spearman and Pearson correlations for the 34 finite gene–timepoint pairs (Tables S63–S64). Lesion staging, microscopy, and fungal: plant DNA biomass qPCR were used only as developmental or biomass context. Transcript RT-qPCR was treated as cross-method concordance for selected source-study transcripts, not independent biological replication or translation validation. The Wei et al. transient-expression screen was used as different-modality heterologous-phenotype evidence for a small source-defined effector subset. Tomato sampling under red light at 23 and 47 HPI was retained as a diel/illumination design caveat. None of these observations were treated as evidence of codon-dependent translation or as a test of the current codon-demand statistic.

A post hoc selected-gene crosswalk joined the 16 Reboledo RT-qPCR genes to the Reboledo putatively secreted set, Wei candidate-effector set, Wei functionally screened candidates, and Wei strong cell-death-inducer subset through reconciled current-reference gene identifiers (Tables S48 and S63). This descriptive overlap was used only to triangulate candidate relevance across source-study evidence types; it was not assigned an inferential probability and was not treated as validation of a complete gene set or codon-demand statistic.

#### SM7. Matched-host analyses

Host codon demand was calculated from the same dual-RNA-seq libraries using each host's own coding sequences and the same coding-length correction. We analyzed the host and fungus separately; we did not share a codon-preference scale or tRNA system between organisms. Host robustness scenarios paralleled the fungal abundance, coding-length, protein-family, and translation-gene exclusions. Complete host robustness, trajectory-reassignment, mapping-QC and host–fungus directional results appear in Figures S9 and S15 and Tables S5 and S67–S69. We compared host and fungal time-aligned 16–24-HPI vectors by cosine similarity in the common codon alphabet but did not compare the magnitudes of species-specific CAI values.

#### SM8. BcNPTII sequence calibration and BcBOT2 context

The published NPTII and synonymous codon-optimized BcNPTII coding sequences encode the same protein and provide an external sequence-calibration example. We calculated their genomic-frequency, ribosomal-reference CAI, balanced transcript-demand, and tRNA-copy proxy percentiles against endogenous B05.10 genes. We explored time-specific relative-abundance potential with study-specific models containing fixed CAI, GC3, and coding length; the response used competitive counts divided by analyzed coding length and closed within the library. Because the construct was designed using codon criteria and neither sequence was expressed in these infection datasets, the analysis is not an independent positive control or observed transgene-expression test. Figure S16 and Tables S70–S71 contain sequence metrics and exploratory predictions.

BcBOT2 is the endogenous deletion target used in the transformation experiment and is not part of the BcNPTII marker coding sequence. Its observed expression was shown only to contextualize the difference between a fixed synonymous marker contrast and an endogenous infection-regulated gene (Table S72).

#### SM9. Decoding, modification-sensitive and local-sequence sensitivity analyses

The genomic B05.10 tRNA reference included 210 predicted nuclear tRNAs and 43 anticodons. We recalculated demand-to-capacity ranks under five fixed decoding schemes spanning mild or strong wobble penalties, with or without an inosine-like proxy. Structural zero-capacity codons were retained as missing states rather than assigned arbitrary small values. Figure S17 and Tables S73–S75 report reference auditing, temporal sensitivity, and rank correlations. These analyses assess dependence on genomic decoding assumptions; they do not measure time-specific tRNA abundance or charging.

Predefined AAA/AAG, CAA/CAG, and GAA/GAG contrasts were tested across all fungal genes and within secreted/extracellular genes against function-conditioned nulls. We tested expression of 15 annotated tRNA-modification machinery candidates across infection timepoints. Figure S18 and Tables S76–S77 contain these results. Transcript abundance of a modifying enzyme was not interpreted as enzyme activity or modification state.

We measured local sequence organization as the mean ribosomal-reference codon-preference residual among the first 30 post-start codons relative to the remainder of each coding sequence. Within-gene synonymous reallocations preserved the encoded protein and the gene's complete codon inventory. Fixed local scores were then weighted by library expression. This component was run with Python 3.12.14, NumPy 2.3.5 and pandas 2.2.3; the saved parameter record also stores these versions. Figure S19 and Tables S78–S80 report fixed sequence patterns, temporal tests, gene scores, and expression coverage.

#### SM10. Lower-replication tomato sensitivity and reproducibility

PRJNA1056687 was processed using the same competitive fungal reference and temporal CLR workflow. Because each timepoint contained two libraries, multivariate dispersion was not estimated: two points have identical distances to their group centroid and provide no reliable residual dispersion. Temporal effect, family-balanced, omission, allocation, and matched-direction results are reported in Figure S10 and Tables S3 and S54. The study was excluded from the primary three-system combined evidence.

All analysis stages used saved parameter files and fixed random seeds. Output manifests recorded the generating script, source file, size, and modification time. Primary statistical and figure generation used R 4.6.0 with data.Table 1.18.4 and ggplot2 4.0.3; mapping-specific R and Rsubread versions are reported in SM2. Empirical probabilities used the plus-one correction, and adjusted probabilities used the Benjamini–Hochberg method within prespecified test families. Table S51 contains the cross-study combined evidence. The supplementary package manifest provides checksums and paths for the complete numbered figure and table series. Complete session records, a machine-readable environment specification, and the public code release identifier must be included in the versioned repository named in the final Data Availability statement.

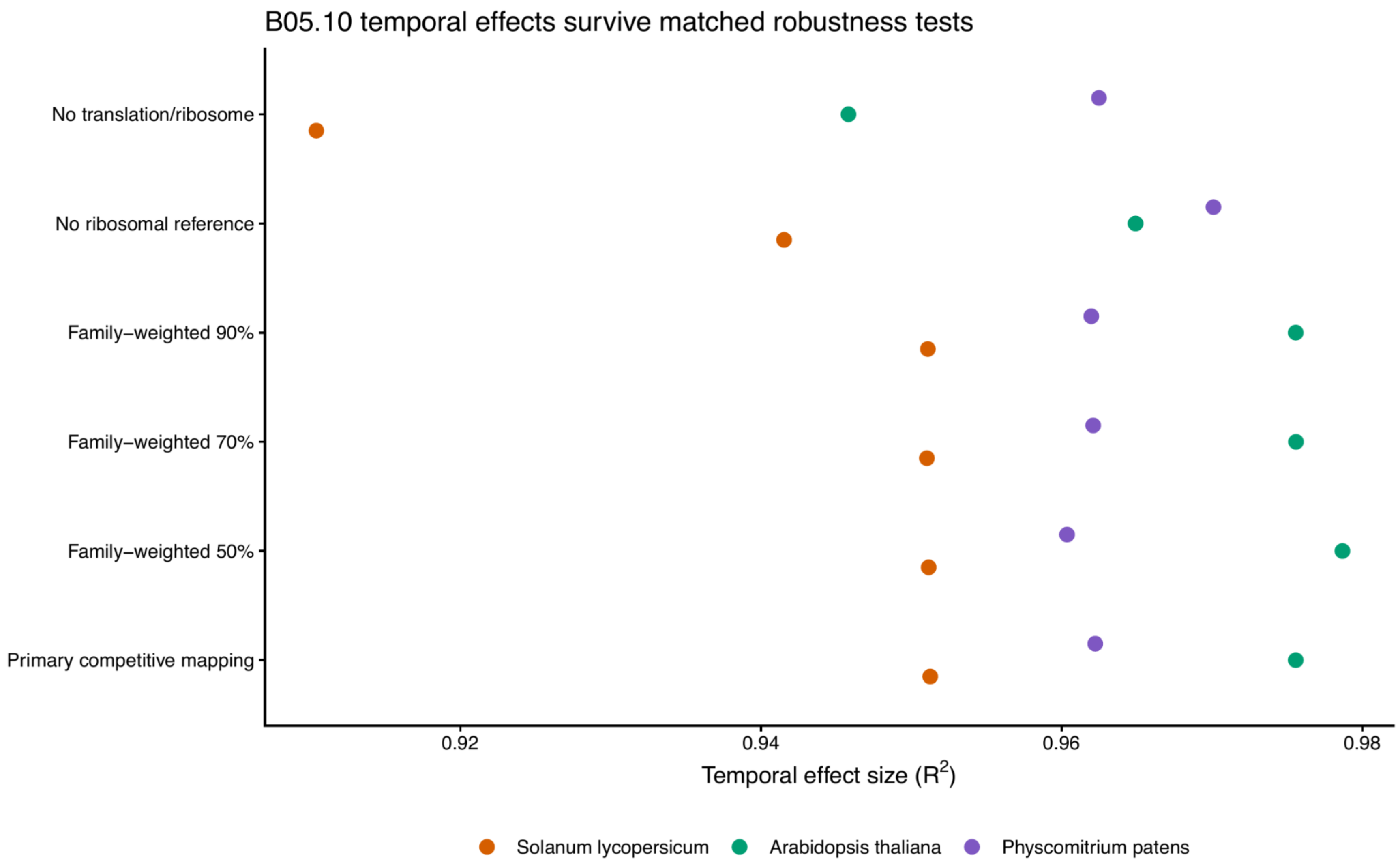


**Figure S1. Robustness to protein-family weighting and translation-gene exclusion**

Study-specific fungal temporal $R^2$ values for the primary length-normalized competitive-mapping analysis, inverse protein-family-size weighting at 50%, 70% and 90% identity, exclusion of the 87-gene ribosomal reference, and exclusion of all annotated ribosome/translation genes. All three primary infection datasets retain significant temporal structure, and sensitivity profiles remain highly correlated with their corresponding primary pattern.

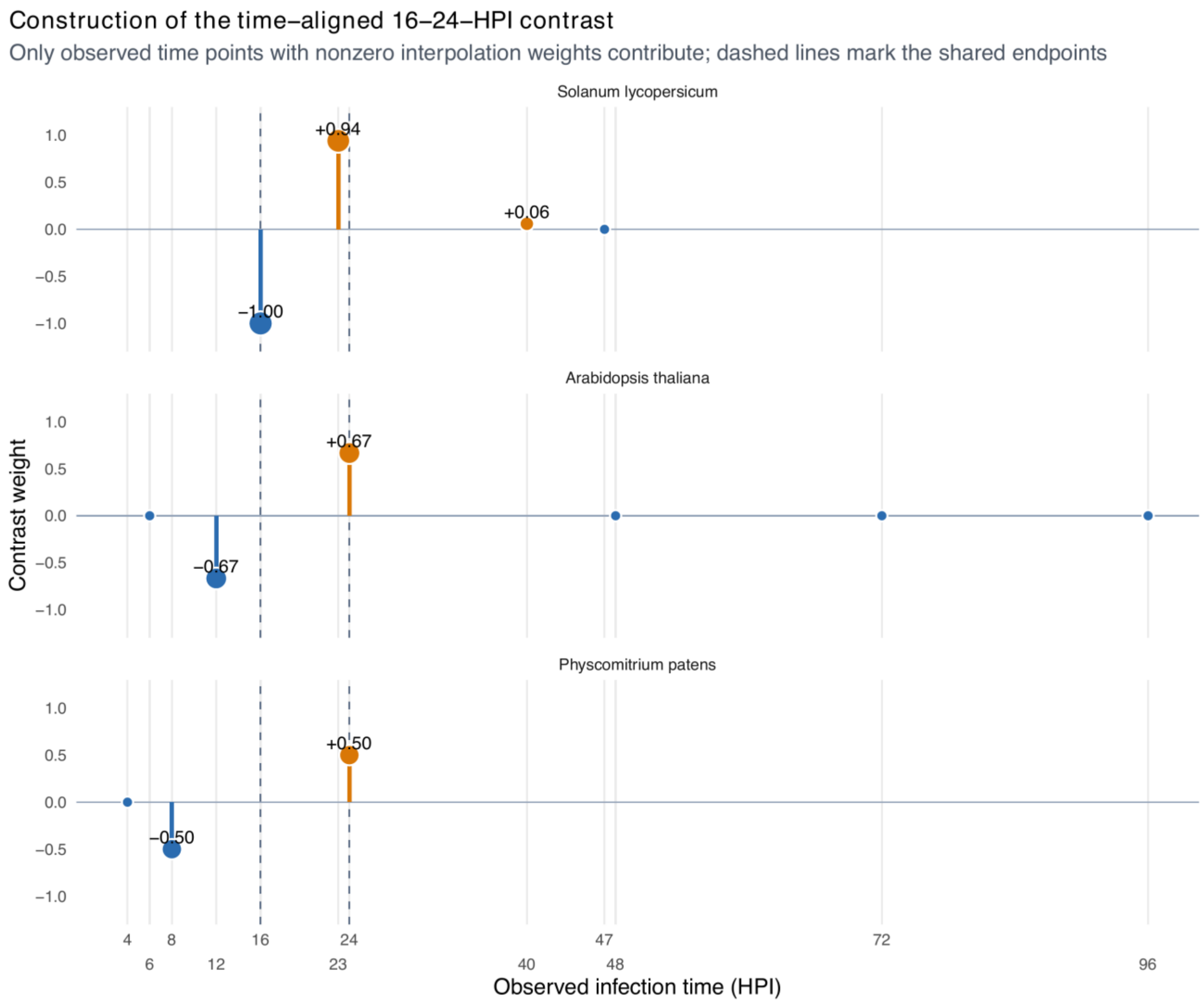


**Figure S2. Construction of the time-aligned 16–24-HPI contrast**

Observed time points and their signed interpolation weights for the common 16–24-HPI fungal contrast. Negative weights reconstruct the 16-HPI endpoint; positive weights reconstruct the 24-HPI endpoint; observed times with zero weight do not contribute. Dashed lines mark the shared HPI endpoints. The figure makes explicit that no dataset observed both endpoints directly and that interpolation used time-point means without creating biological replicates or asserting equivalent developmental state.

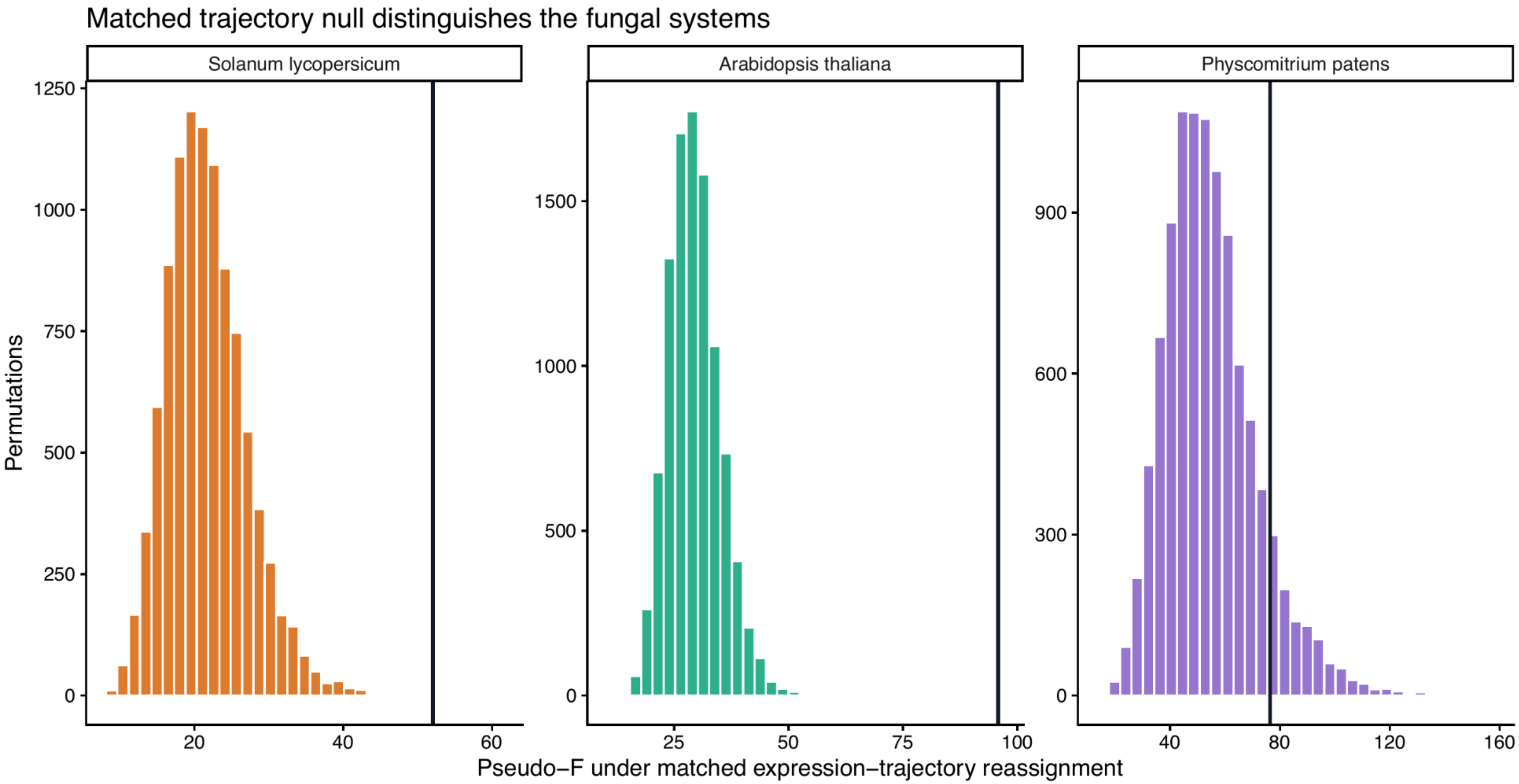


**Figure S3. Matched expression-trajectory reassignment null**

Study-specific null distributions of fungal full-series pseudo-*F* statistics after complete expression trajectories were reassigned among genes matched for mean expression, CDS length, GC3, and amino-acid composition. The observed effect exceeds the 9,999-permutation null in *Solanum lycopersicum* (empirical $P = 0.0004$) and *Arabidopsis thaliana* ($P = 0.0001$), but not in the shorter three-timepoint *Physcomitrium patens* series ($P = 0.1015$). *Physcomitrium patens* nevertheless retains a strong temporal effect in the primary and sensitivity analyses. Figure S4 tests the shared early window and qualifies the full-series interpretation.

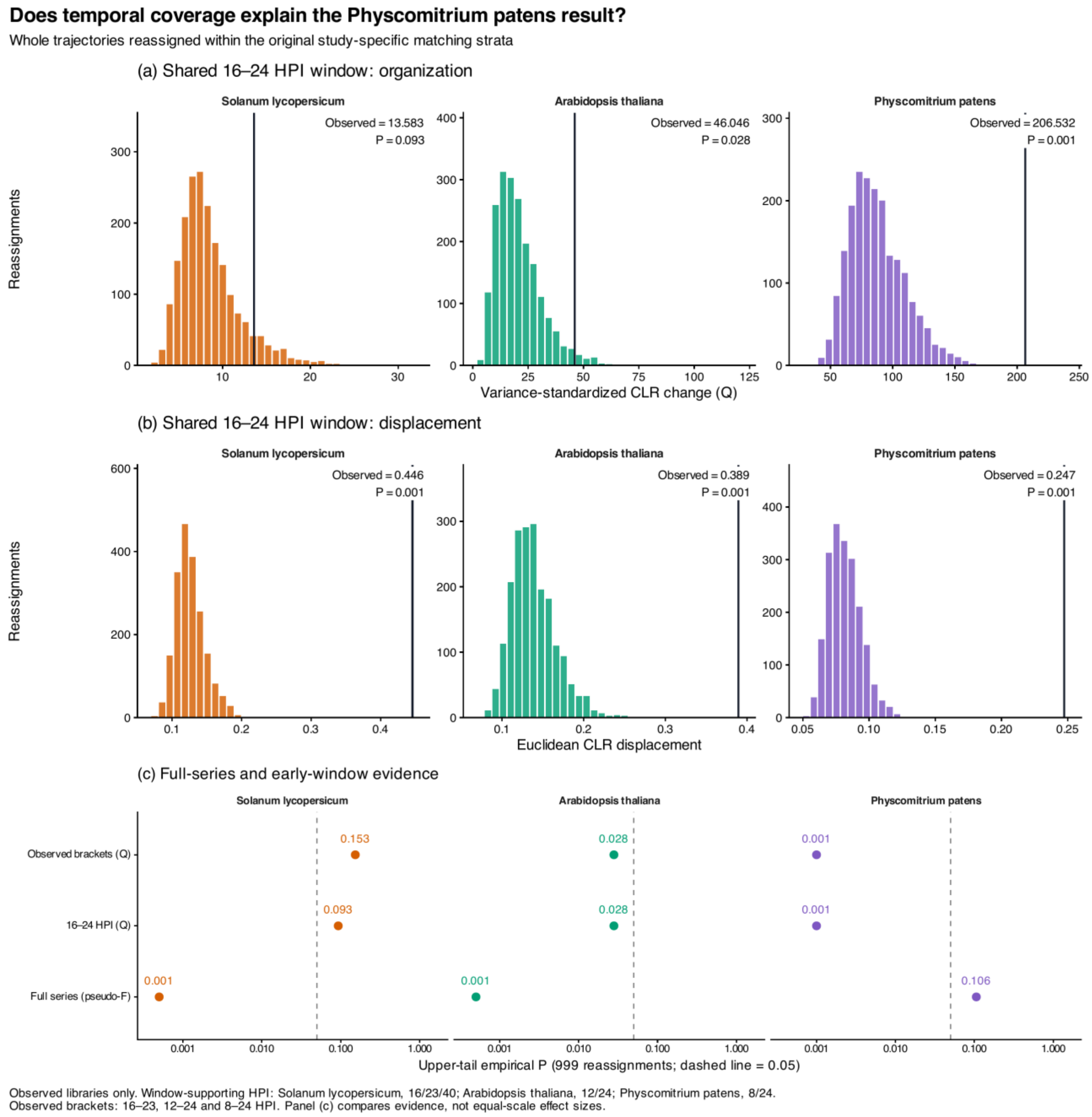


**Figure S4. Shared early-window codon-demand organization and displacement**

**(a)** Null distributions of trace-standardized 16–24 HPI CLR change (Q) for *Solanum lycopersicum*, *Arabidopsis thaliana*, and *Physcomitrium patens*. **(b)** Null distributions of Euclidean CLR displacement over the same contrast. Colored histograms show 1,999 whole-gene trajectory reassignments within the original study-specific strata matched on mean expression, CDS length, GC3, and two amino-acid-composition PCs. Black lines mark the observed statistics; labels give unadjusted upper-tail empirical P. Panel (a) gives $P = 0.093$, 0.028, and 0.001 (BH $q = 0.093$, 0.042, and 0.003); panel (b) gives $P = q = 0.0005$ for each system. Facets have independent axes. **(c)** Full-series pseudo-F, early-window Q, and observed-bracket Q evidence from the same assignments. The dashed line marks unadjusted P = 0.05; probabilities compare evidence, not equal-scale effect sizes or formal between-host differences. Full-series $P = 0.0005$, 0.0005, and 0.106 in this 1,999-permutation shared-window rerun; Figure S3 reports the higher-resolution 9,999-permutation full-series analysis.

All variance estimates use original triplicate libraries. The contrast weights at observed HPI are (−1, 16/17, 1/17) at *Solanum lycopersicum*'s 16/23/40 HPI, (−2/3, 2/3) at *Arabidopsis thaliana* 12/24 HPI, and (−1/2, 1/2) at *Physcomitrium patens* 8/24 HPI. Interpolation defines the interval; it creates no replicate observations. Observed-bracket comparisons are 16–23, 12–24, and 8–24 HPI ($P = 0.153$, 0.028, and 0.001). For the latter two systems, Q is invariant to interpolation scaling and cannot localize movement within the observed interval. Rebuilding expression-matching bins from supporting timepoints gives Q-test $P = 0.074$, 0.0245, and 0.0005. See Tables S43–S45 for full results and sampling provenance.

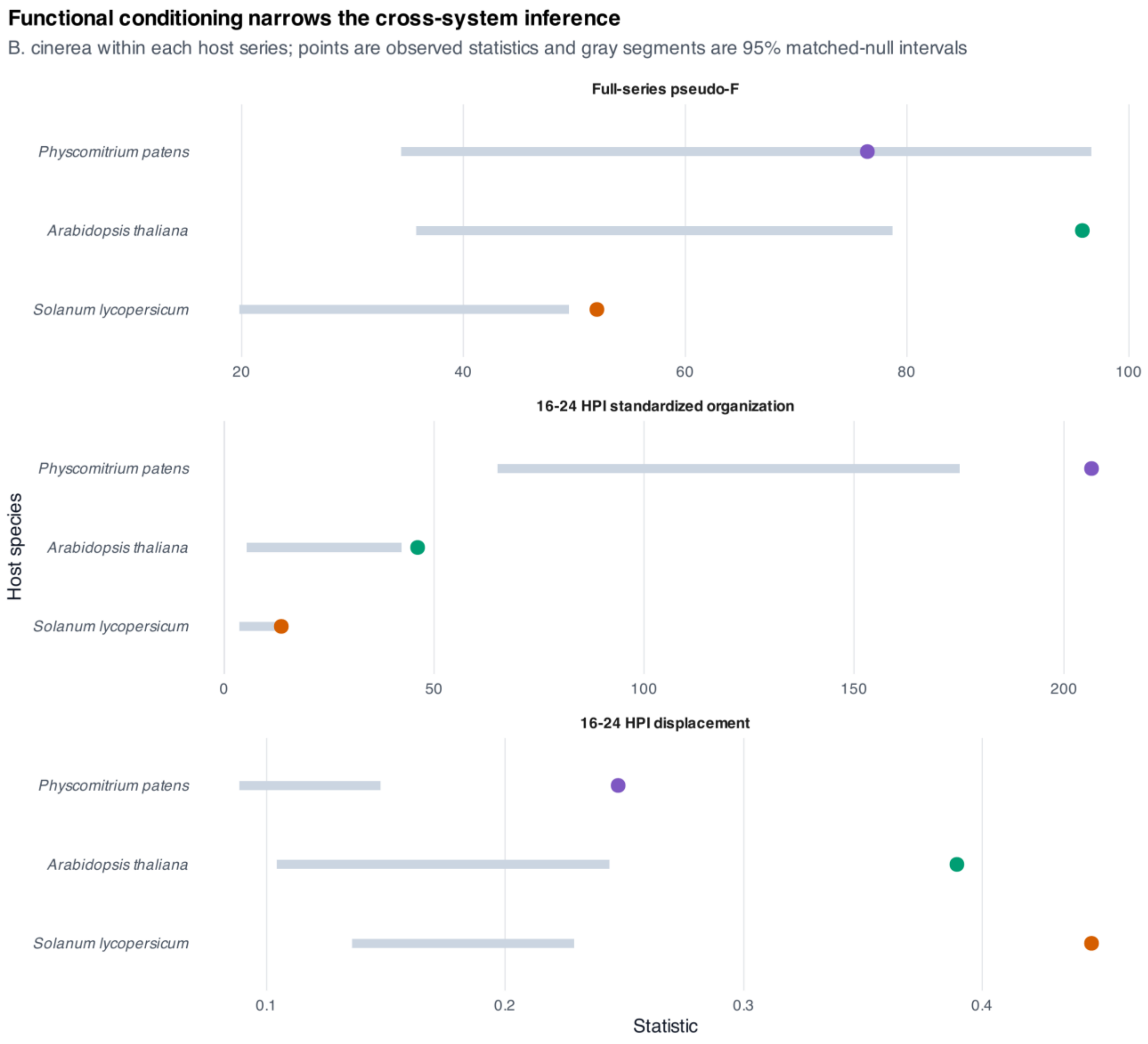


**Figure S5. Functional-group-conditioned trajectory reassignment**

Observed full-series pseudo-*F*, trace-standardized 16–24 HPI organization, and 16–24 HPI CLR displacement are shown as colored points; gray segments show the 95% intervals from 1,999 whole-expression-trajectory reassignments restricted by broad UniProt-derived functional group, mean expression, CDS length, GC3, and two amino-acid-composition PCs. More than 99.4% of genes remain movable in every system. Full-series excess persists in *Solanum lycopersicum* and *Arabidopsis thaliana* but not *Physcomitrium patens*. Shared-window displacement exceeds the null in all systems (BH $q = 0.0005$ each), as does trace-standardized organization (BH $q = 0.018$ each).

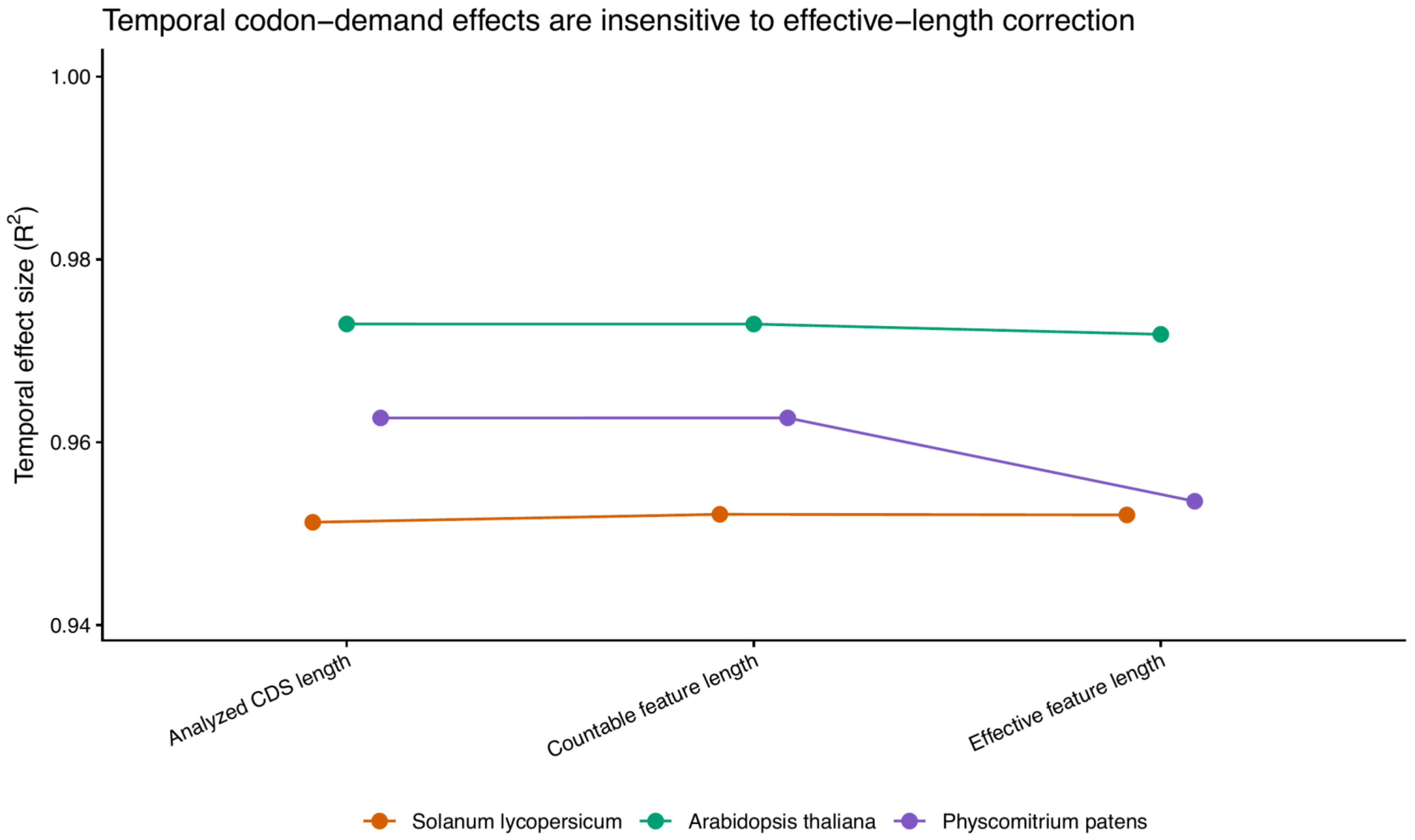


**Figure S6. Effective-length sensitivity of temporal codon demand**

Temporal multivariate effect size ($R^2$) under the primary analyzed-CDS length, the exact annotation feature length supplied to counting, and a library-specific effective feature length equal to feature length minus mean fragment or read length plus one. The same eligible genes and fixed competitive count matrices were used for all three definitions. The sensitivity addresses countable-feature and fragment-end opportunity but is not isoform-level expectation maximization or sequence-, positional-, or GC-bias correction.

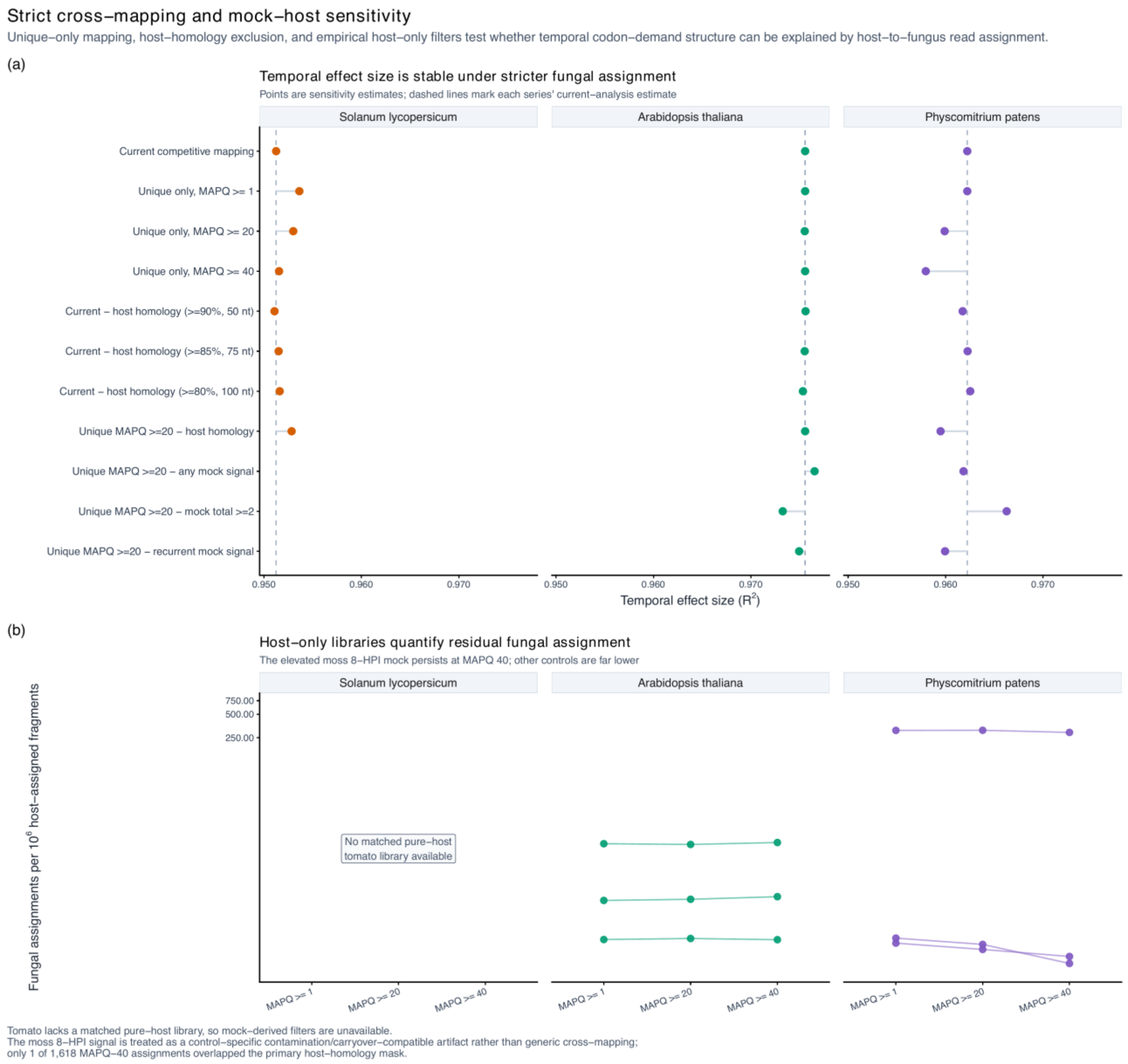


**Figure S7. Temporal inference is tested under strict cross-mapping and mock-host controls**

**(a)** Within-study temporal $R^2$ after current competitive assignment, unique-only competitive transcript mapping at MAPQ thresholds 1, 20 and 40, conservative removal of B05.10 genes with host-similar sequence, and—in the systems with pure host controls—removal of fungal genes receiving signal in a host-only library. Points are separate sensitivity analyses; the primary host-homology threshold was ≥90% nucleotide identity over ≥50 nt, and the primary empirical blacklist removed any gene with a MAPQ≥20 fungal count in a selected host-only library. The current-analysis estimate is shown as the reference within each system. **(b)** False fungal assignment in three uninfected *Arabidopsis* libraries and three mock-inoculated moss libraries, expressed per million host-assigned fragments across MAPQ thresholds. The primary tomato series has no pure host-only deposited control because its 0-HPI libraries are host-plus-spore synthetic mixtures. These controls estimate assignment leakage under the selected reference and mapping rules; they do not prove the origin of every infection read. One moss 8-HPI mock had an isolated, MAPQ-resistant apparent fungal signal, whereas its 4- and 24-HPI mocks were below one fungal assignment per million host assignments. It is treated as compatible with control-specific contamination or carryover rather than generic sequence cross-mapping because only one of its 1,618 MAPQ≥40 assignments overlapped the primary host-homology mask. Any-signal, total-count, and recurrent-control blacklists are reported separately; Table S52 gives the full control-by-mask overlap audit.

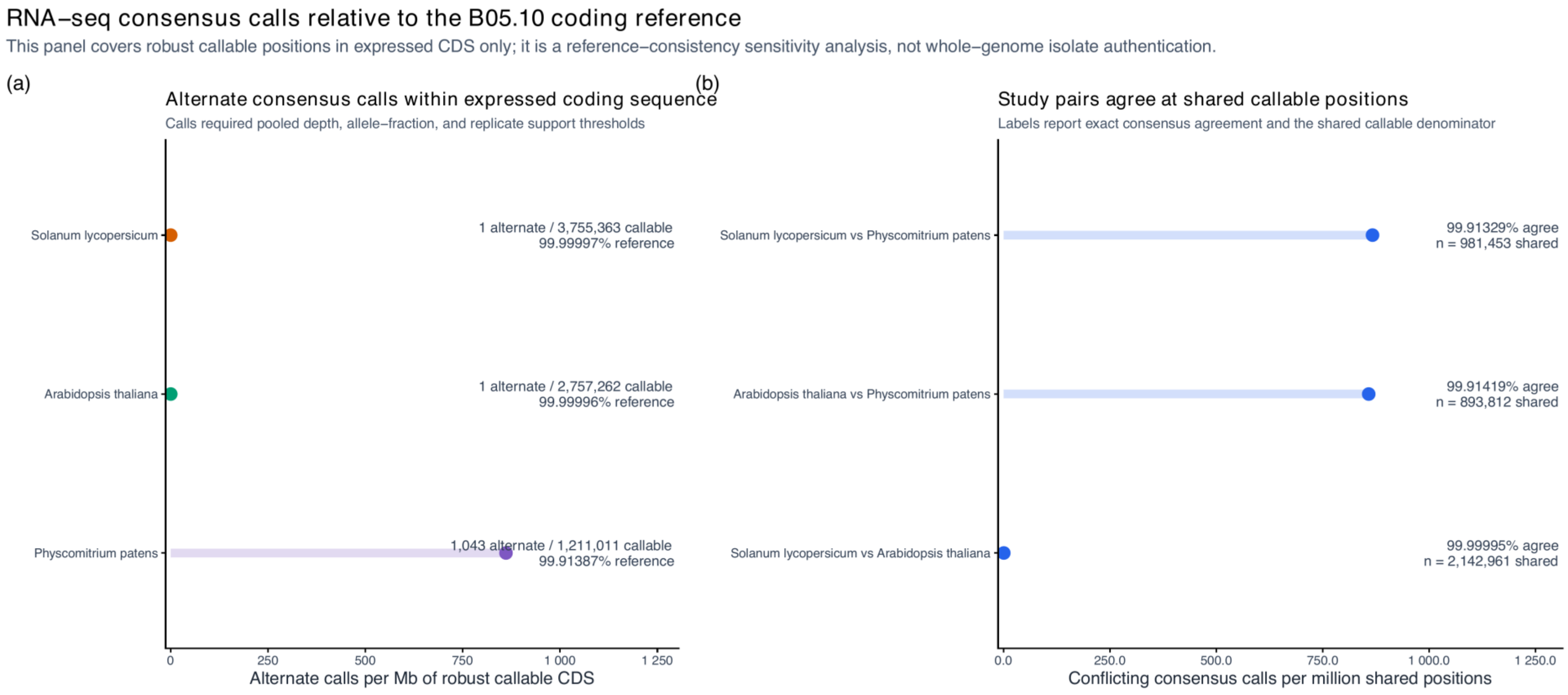


**Figure S8. Expressed coding sites quantify reference concordance without authenticating isolate identity**

**(a)** High-confidence nonreference consensus calls per million robustly callable B05.10 coding positions in three predeclared high-fungal-depth libraries from each study. Labels report reference concordance and callable positions. **(b)** Pairwise consensus discordance per million coding positions callable in both studies; labels report the number of shared callable sites. Candidate sites required unique competitive mapping, MAPQ ≥20, base quality ≥30, pooled depth ≥30, major-allele fraction ≥0.90, and support in at least two libraries with per-library depth ≥5 and major-allele fraction ≥0.80. Alignments allowed at most six mismatches, and the first and last ten bases of every CDS were excluded. The panel samples represent coding sequences and are not whole-genome isolates; mapping to the B05.10 reference may also favor the reference allele.

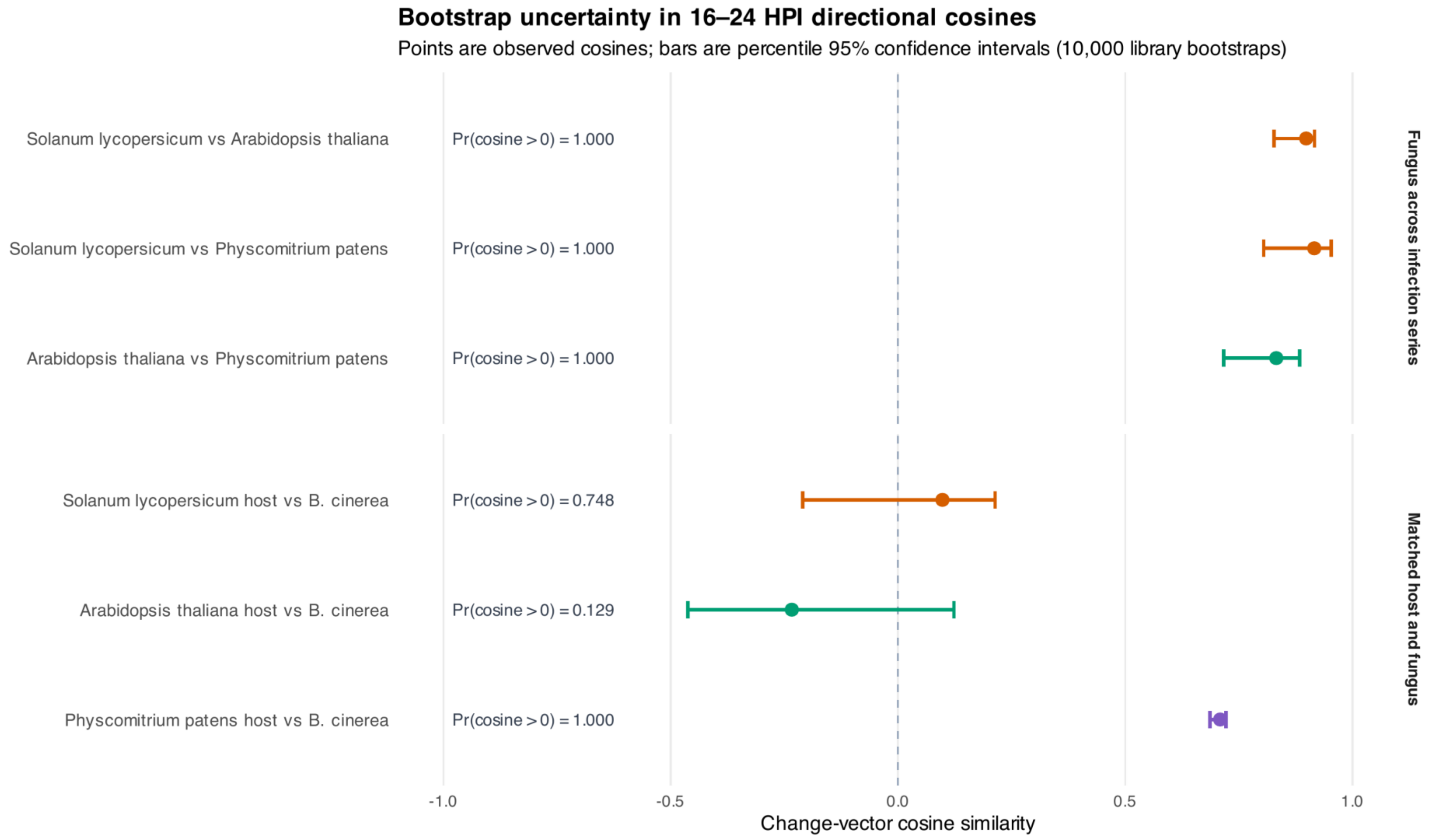


**Figure S9. Bootstrap uncertainty in time-aligned directional cosines**

Observed change-vector cosine similarities and 95% percentile confidence intervals from 10,000 resamplings of biological libraries within each observed time point. The upper panel compares fungal 16–24-HPI vectors among infection series; the lower panel compares host and fungus within each series. Timepoint centroids were linearly interpolated only within observed HPI ranges after resampling. Labels report the bootstrap fraction with cosine greater than zero; this is a directional stability summary, not a formal test of host causality.

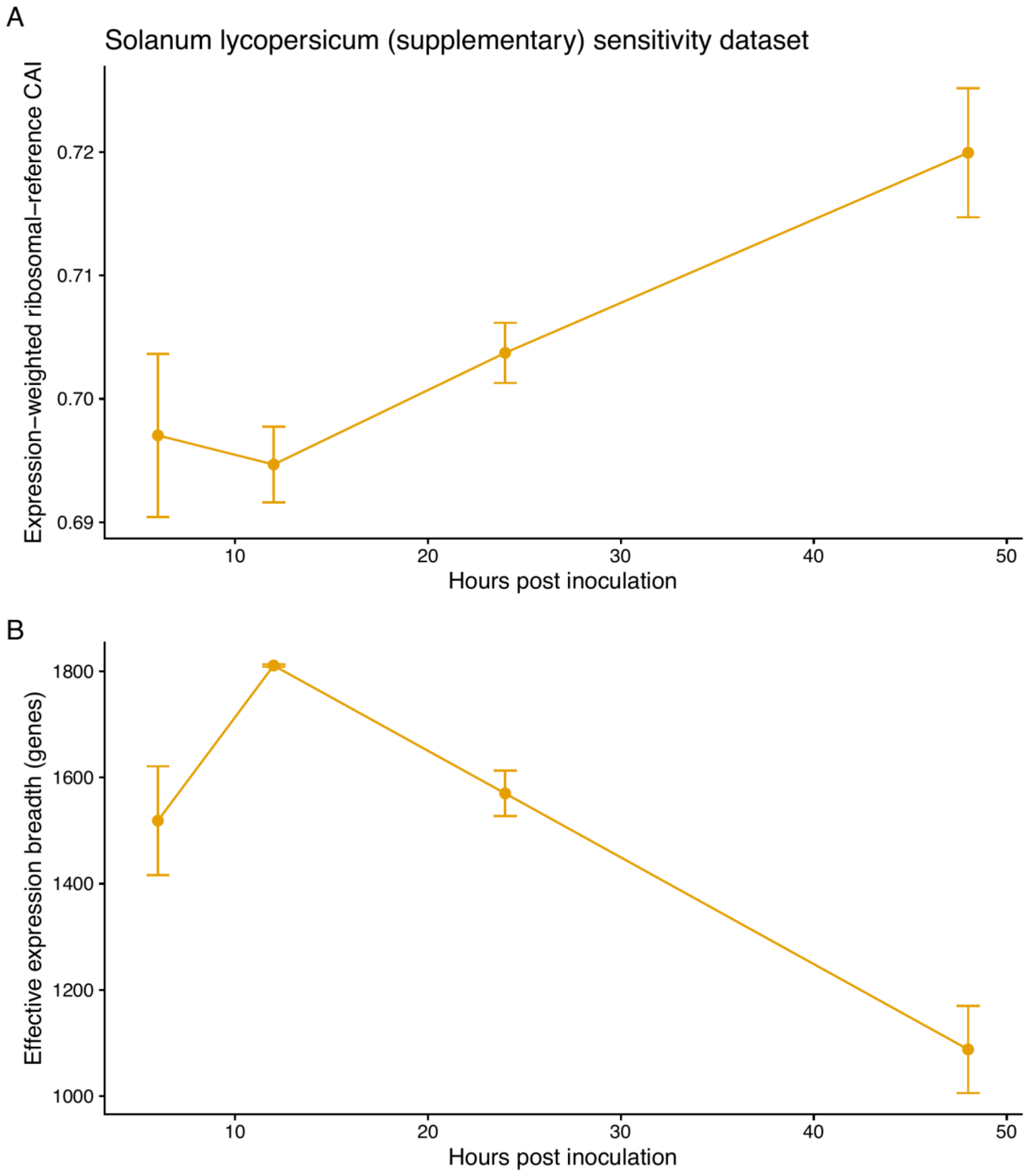


**Figure S10. *Solanum lycopersicum* (supplementary) sensitivity analysis**

Timepoint means ± SE for length-corrected, abundance-weighted ribosomal-reference CAI and effective expression breadth in the lower-replication *Solanum lycopersicum* sensitivity dataset (PRJNA1056687). The overall temporal point estimate is large ($R^2 = 0.866$) but does not pass the 9,999-permutation test ($P = 0.0561$); expression breadth varies by time ($P = 0.0096$), whereas weighted CAI does not ($P = 0.0579$). This series is reported separately because two libraries per timepoint preclude a reliable dispersion test; its result is less stable to library omission and CV. Moneymaker reads were mapped against the available M82 host transcript reference.

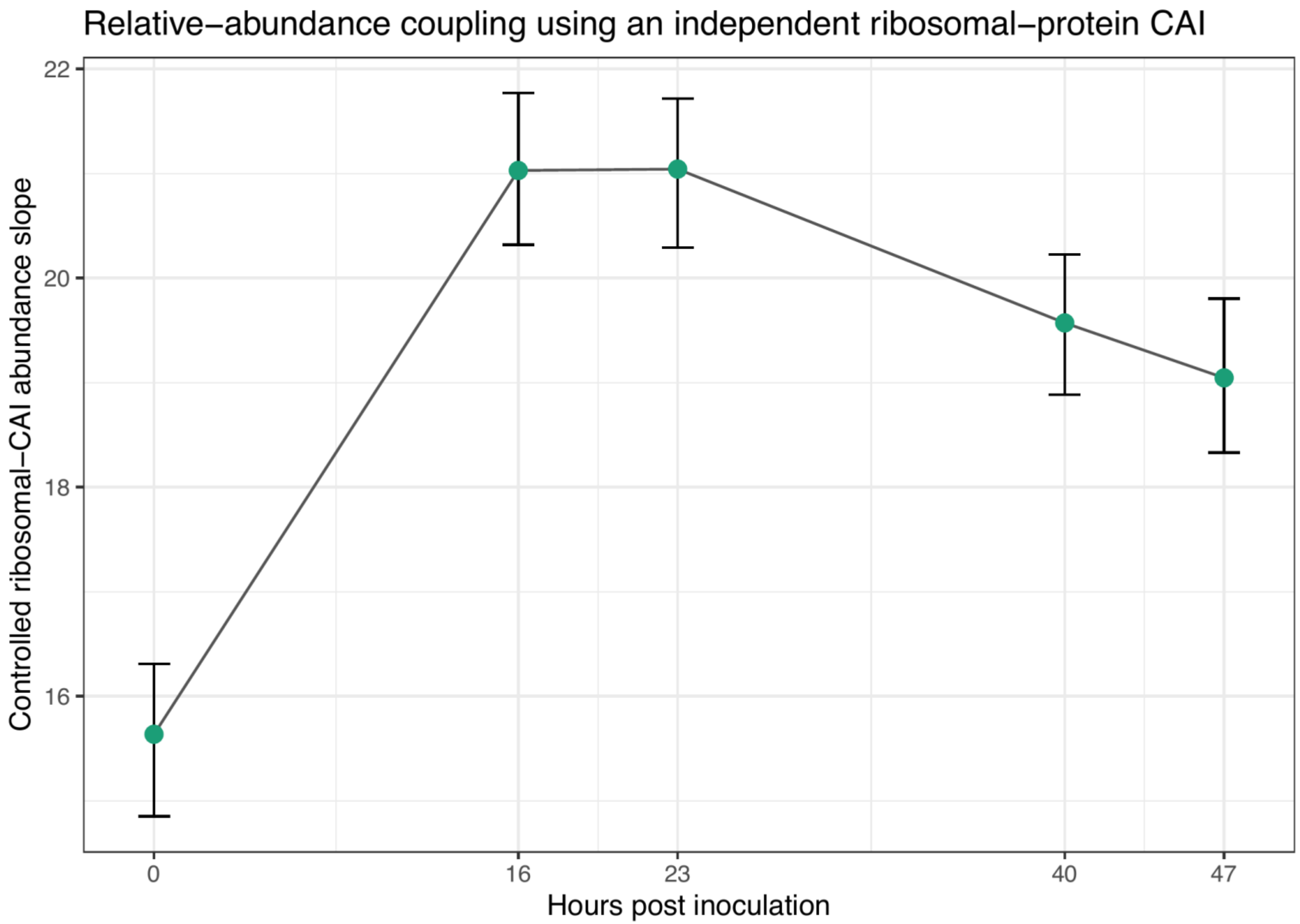


**Figure S11. Independent ribosomal-reference CAI analysis**

Controlled relative-transcript-abundance slopes calculated using fixed CAI weights derived from 87 annotated ribosomal-protein genes. Competitive counts were divided by analyzed coding length before library closure. Positive slopes at every infection timepoint show that higher-CAI genes tend to have greater relative abundance after adjustment for GC3 and protein length; the weaker slopes at 40 and 47 HPI complement the decline in abundance-weighted CAI.

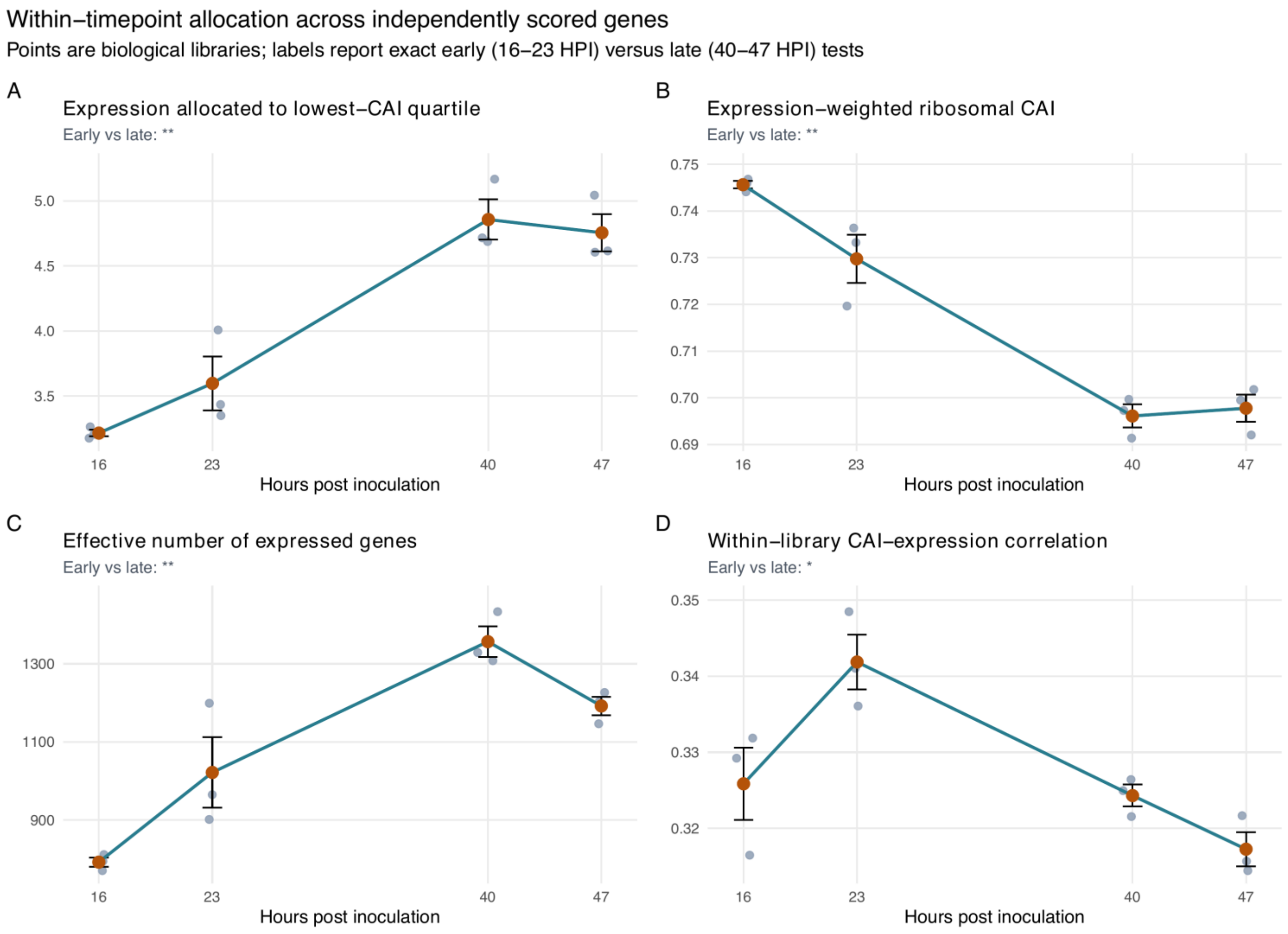


**Figure S12. Tomato-series detailed transcript-allocation metrics**

Library values and timepoint mean ± SE for abundance assigned to the lowest-CAI quartile, expression-weighted CAI, effective expression breadth, and the within-library CAI-expression correlation. Panel subtitles report exact 16–23-versus-40–47 HPI tests corrected across allocation metrics. Main Figure 5 places the two directly comparable allocation summaries across all three host series.

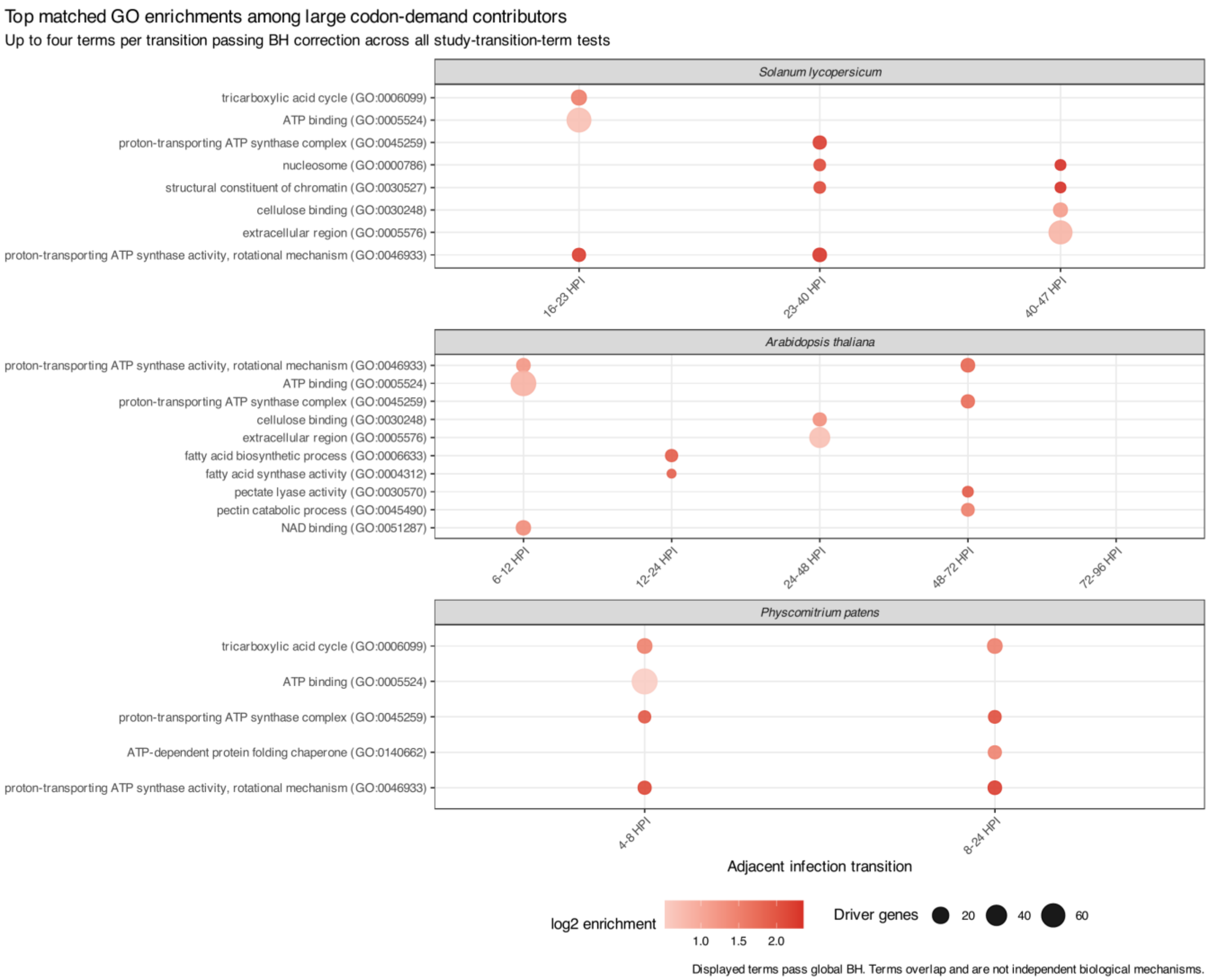


**Figure S13. Non-translation programs reshape synonymous codon demand**

GO enrichment among the top 5% of recalculated temporal driver genes after excluding 456 genes annotated for ribosome- or translation-related functions. Enrichment is evaluated against 4,999 null gene sets matched to the observed drivers within coarsened joint strata of study-wide mean relative transcript-molecule abundance, analyzed CDS length, and transition-specific abundance-change magnitude. Panels show up to four terms per adjacent transition that pass Benjamini–Hochberg correction across all 8,890 study-transition-term tests; the Arabidopsis 72–96-HPI transition has no term passing that threshold. Point size indicates the number of selected driver genes annotated to the term, and color indicates the log2 matched enrichment ratio. Globally corrected enrichments occurred in nine of ten transitions and comprised 37 term-transition associations representing 16 distinct GO terms. These are overlapping annotations among genes that disproportionately shape the RNA-level codon-demand trajectory, not independent mechanisms, pathway activity measurements, or evidence of translation efficiency.

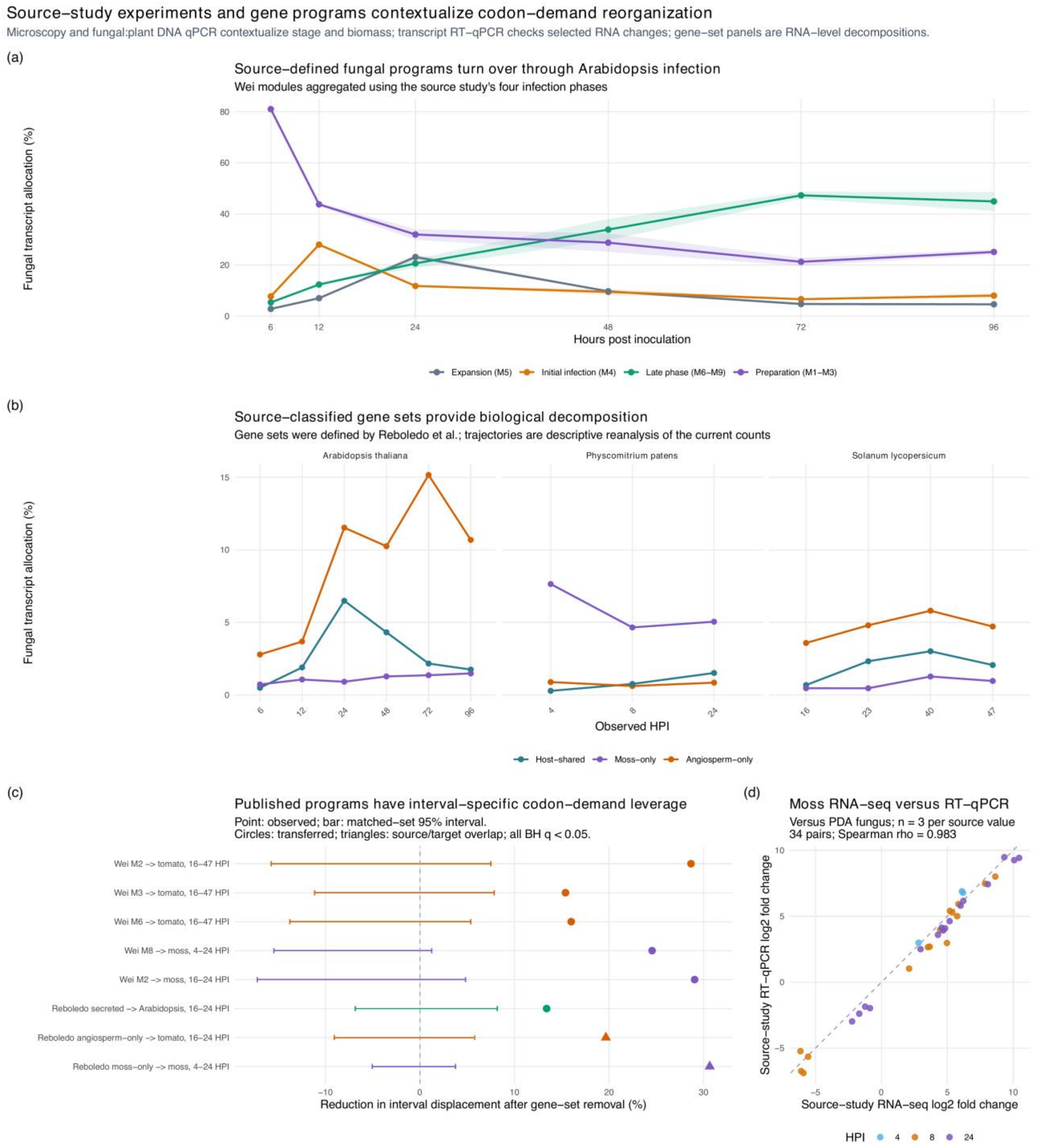


**Figure S14. Source-study experiments and gene programs contextualize codon-demand reorganization**

**(a)** Relative fungal transcript-molecule allocation to the nine *Arabidopsis* co-expression modules reported by Wei et al., aggregated into their four source-defined infection phases. Lines show timepoint means and ribbons show standard errors among three biological libraries. **(b)** Allocation to the host-shared, moss-only, and angiosperm-only upregulated gene sets reported by Reboledo et al., recalculated from the current competitive counts. **(c)** Reduction in the labeled interval's codon-demand displacement after selected source-defined gene sets were removed. Circles identify sets transferred into an RNA-seq series other than the one used to define them; triangles identify source and target RNA-seq that overlap. Points are observed leave-set-out attenuation, and bars are 95% intervals from the 999-draw screening matcher, which prioritized the same abundance-, coding-length-, and GC3 cell but expanded the candidate pool when exact-cell capacity was insufficient. The panel displays selected significant transferred examples and two significant source/target-overlap examples; Table S66 reports every tested source set and contrast. All displayed results have full-family BH-adjusted $P < 0.05$ in the screen. A separate strict sensitivity preserves exact cell counts and balances interval-specific RNA-allocation change magnitude and amino-acid-composition PC1–PC2; it does not match signed change or within-set coordination. Attenuation is interpreted as disproportionate contribution under the stated matching model, not unique codon-compositional leverage. **(d)** Cross-method concordance between RNA-seq and transcript RT-qPCR log2 fold changes transcribed from the Reboledo source study for 34 finite gene–timepoint pairs among 16 fungal pathogenesis genes. Both assays reported fold change relative to PDA-grown fungus, and each source value summarized three biological replicates. The source paper reported regression $R^2 = 0.9882$; the displayed pooled Spearman *rho* is a new descriptive recalculation. The dashed line is equality. Microscopy, fungal: plant DNA biomass qPCR, transcript RT-qPCR and heterologous phenotypes are cataloged in Table S62; only the transcript RT-qPCR comparison is plotted. The RT-qPCR values check selected source-study transcripts across methods but are not independent biological replicates. Microscopy and biomass measurements support developmental staging, while module and gene-set analyses decompose partly shared RNA-seq data. None validates the codon-demand statistic or translation.

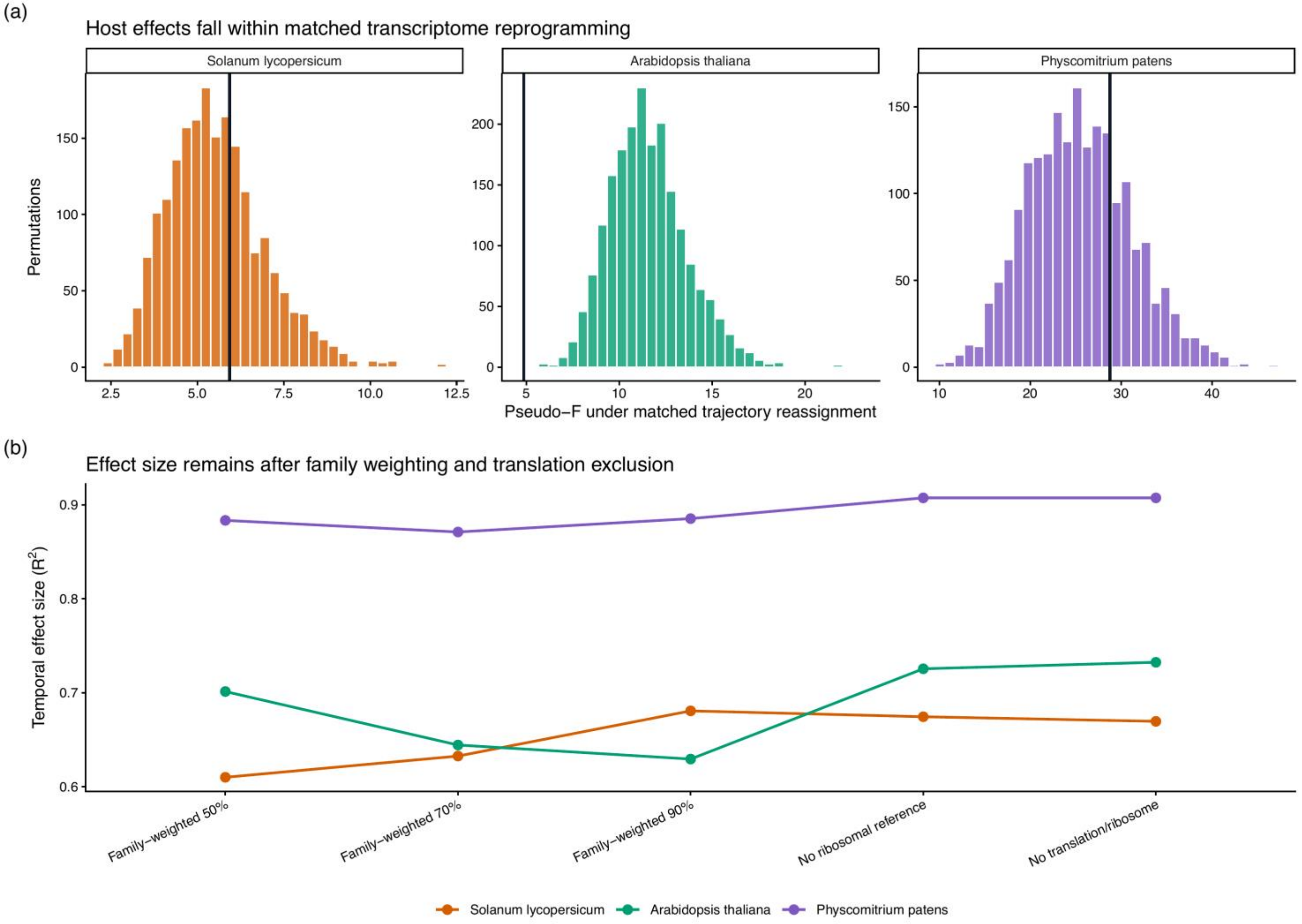


**Figure S15. Matched host robustness and trajectory nulls**

**(a)** Matched whole-trajectory reassignment nulls for the three hosts; black vertical lines mark the observed temporal pseudo-*F*. None of the host temporal effects exceed their matched nulls (*Solanum lycopersicum* empirical $P = 0.350$; *Arabidopsis thaliana*, 1.000; *Physcomitrium patens*, 0.269). **(b)** Study-specific host temporal pseudo-$R^2$ values remain similar after protein-family weighting and exclusion of ribosomal-reference or broader translation-associated genes.

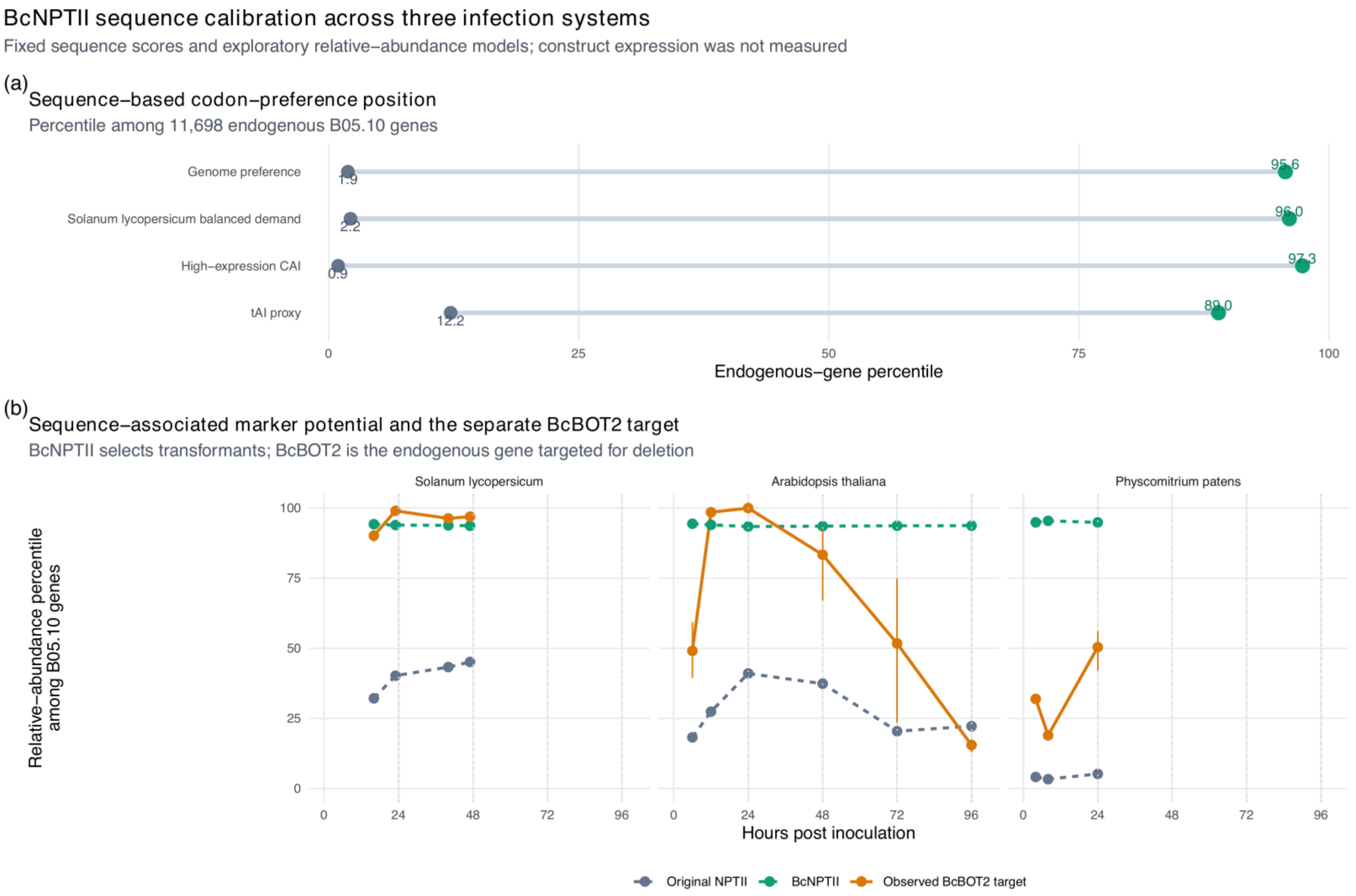


**Figure S16. BcNPTII external sequence-calibration example**

NPTII/BcNPTII is the selectable-marker gene; BcBOT2 is a separate endogenous deletion target. In the published pBOT2-ko vector, the BcNPTII cassette enables selection while BOT2-targeting flanks direct disruption of BcBOT2. BcBOT2 is therefore not part of the marker coding sequence.

**(a)** Percentile positions of the original NPTII and codon-optimized BcNPTII coding sequences relative to endogenous B05.10 genes. Connected points emphasize the paired change for four codon-preference metrics. **(b)** Sequence-associated relative-transcript-abundance potential for the original NPTII and BcNPTII sequences at each infection time point from models fitted independently in *Solanum lycopersicum*, *Arabidopsis thaliana*, and *Physcomitrium patens* using CAI, GC3, and protein length. Model responses use competitive counts divided by analyzed coding length and closed within the library. Observed relative abundance of the endogenous knockout target BcBOT2 (BCIN_12g06390) is overlaid as a biological comparator. Values are percentiles among endogenous B05.10 genes; error bars for BcBOT2 span biological libraries. Across systems, original NPTII occupies approximately the 3.3rd–45.1st percentiles and BcNPTII the 93.4th–95.4th percentiles. Cross-validated R-squared values of 0.10–0.23 show that coding sequence explains only a minority of endogenous expression variation. Neither construct occurs in these RNA-seq datasets, so the marker lines are calibrated to sequence-based expectations rather than observed transgene expression. The published experiment measured transformation and G418-resistance phenotypes, not NPTII transcript abundance.

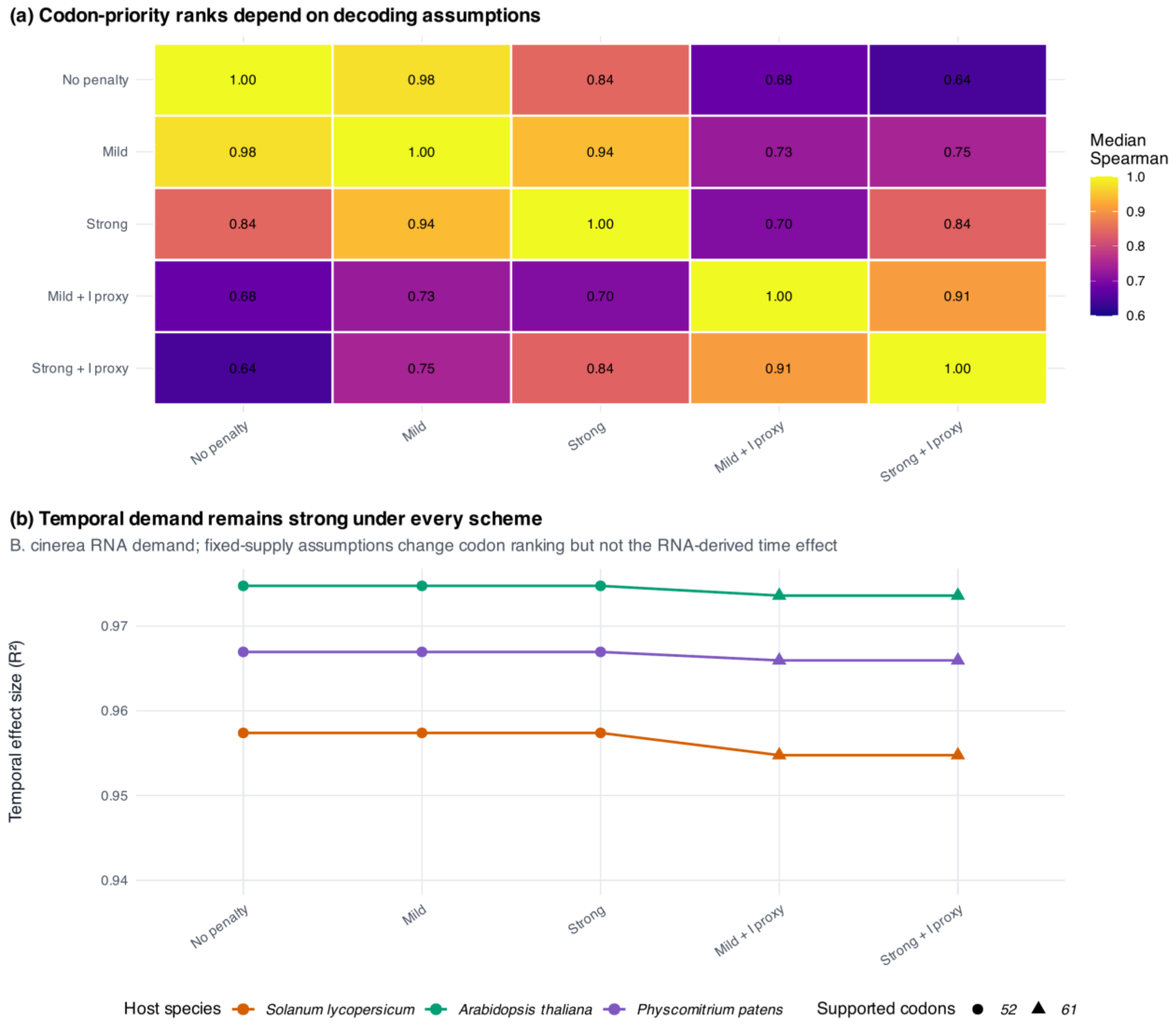


**Figure S17. Sensitivity to genomic tRNA decoding assumptions**

**(a)** Median Spearman correlation of codon demand-to-capacity ranks across 39 fungal libraries under five fixed B05.10 genomic decoding schemes. Comparisons with versus without an inosine-like proxy show the greatest rank sensitivity. **(b)** Temporal $R^2$ for each study and scheme. Circles denote the 52 codons with nonzero capacity without an inosine proxy; triangles denote schemes supporting all 61 sense codons. Every temporal test remains significant. Because the genomic tRNA supply is fixed, these results test sensitivity of codon prioritization and coordinate inclusion, not dynamic tRNA availability.

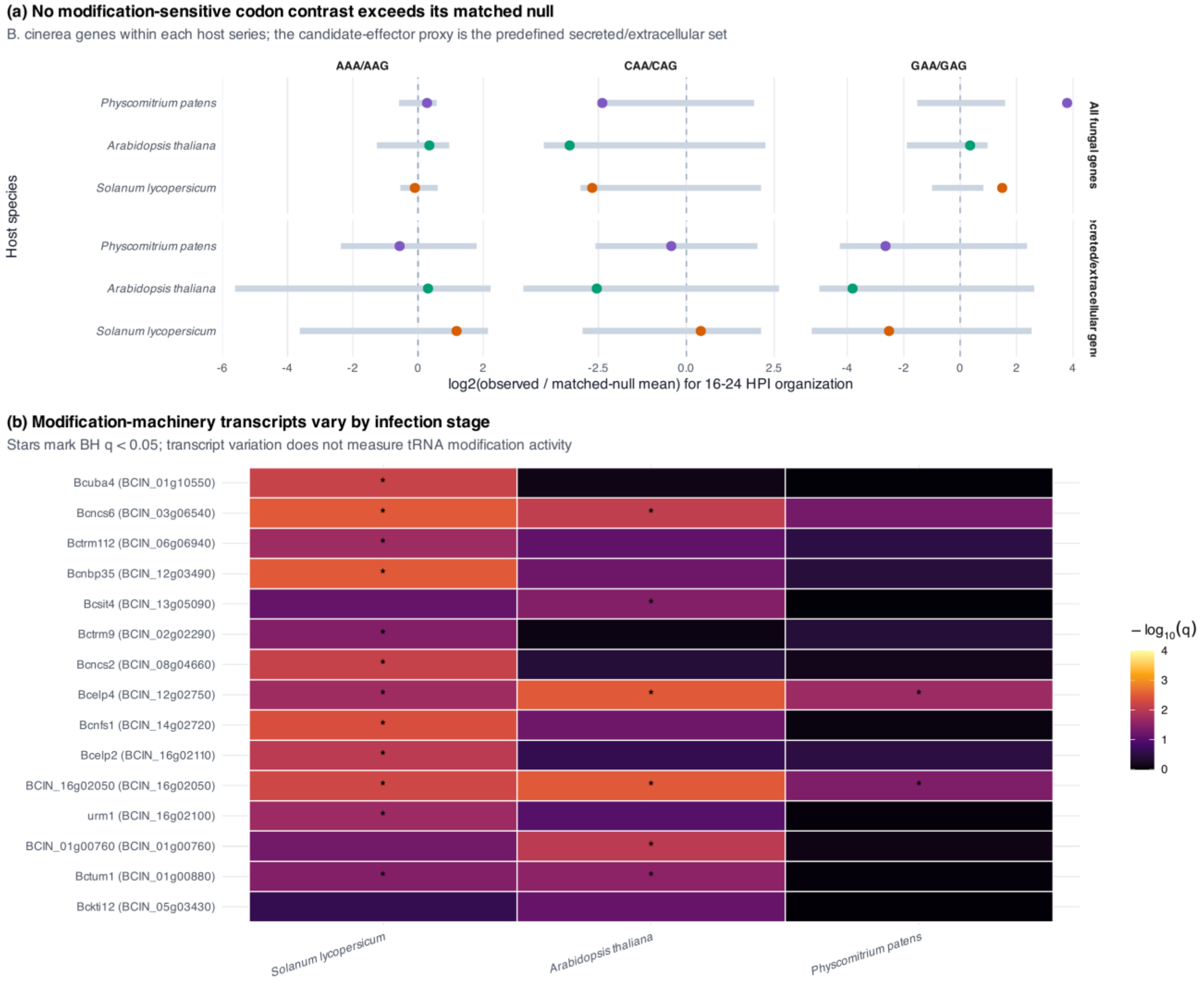


**Figure S18. Predefined modification-sensitive codon and machinery tests**

**(a)** Observed 16–24 HPI organization of AAA/AAG, CAA/CAG, and GAA/GAG log-demand contrasts relative to the function-conditioned matched-null mean. Gray segments show null 95% intervals. Tests were run across all fungal genes and within the predefined secreted/extracellular candidate set. The all-gene GAA/GAG contrast exceeds the null in *Solanum lycopersicum* and *Physcomitrium patens* (BH $q = 0.0045$ each), but not *Arabidopsis thaliana*; no contrast passes correction within the secreted/extracellular set. **(b)** Time-label permutation evidence for time-dependent RNA abundance among 15 B05.10 genes annotated for wobble-uridine modification or tRNA thiolation. Stars mark BH $q < 0.05$. Transcript variation does not measure enzyme activity or tRNA modification state.

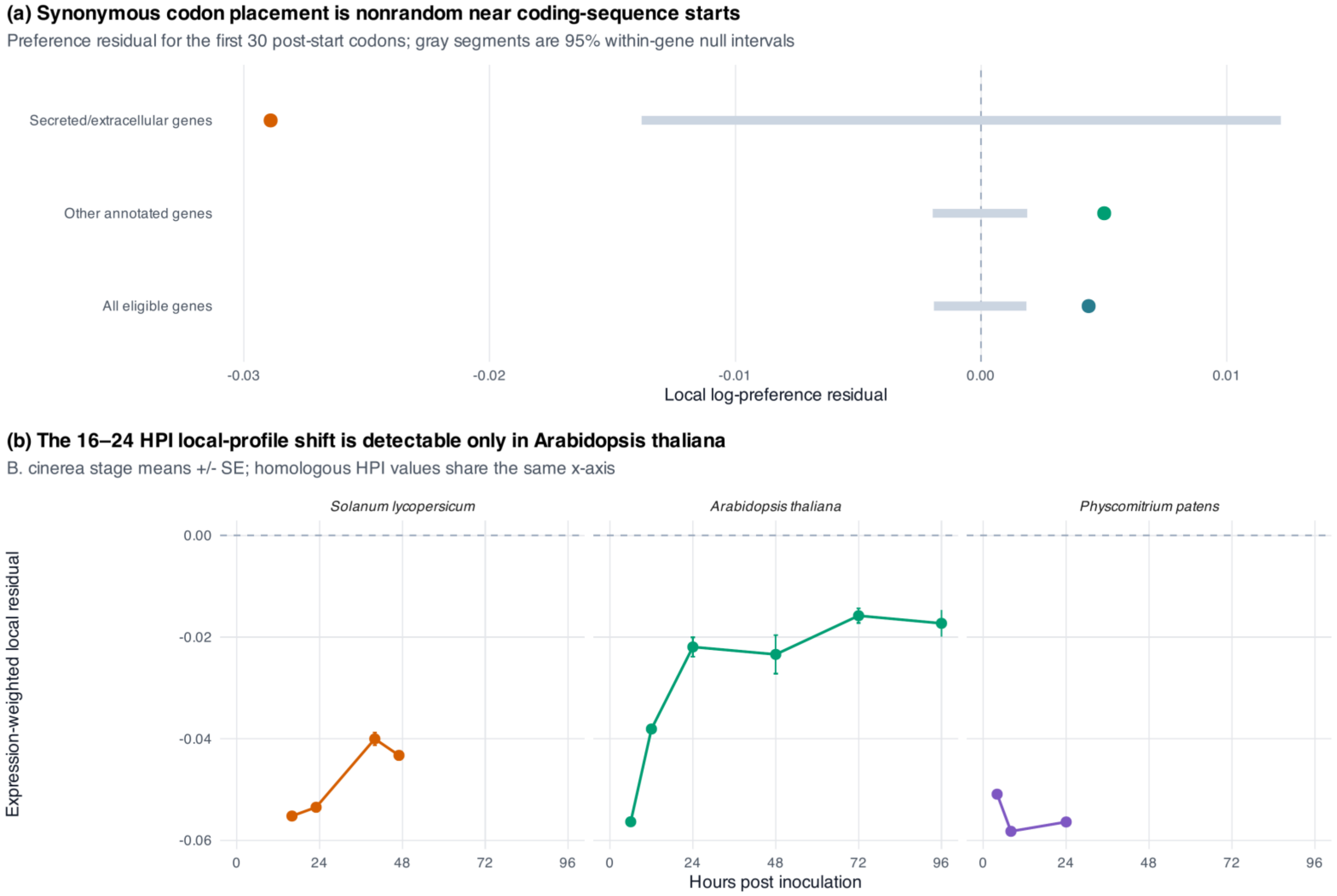


**Figure S19. Local synonymous placement near coding-sequence starts**

**(a)** Mean log ribosomal-reference codon-preference residual in the first 30 post-start codons relative to 999 within-gene synonymous reallocations that preserve protein sequence and each gene's complete codon inventory. Points are observed means, and gray segments are null 95% intervals. Eligible secreted/extracellular genes show a negative local residual, whereas all other annotated genes show small positive residuals. **(b)** Timepoint mean ± SE of the expression-weighted fixed local score, plotted on a common 0–96 HPI axis. Only the *Arabidopsis thaliana* 16–24 HPI absolute change exceeds its local null after correction. These are sequence-context statistics, not measurements of translation initiation, folding, or secretion.
